# Restoring Oscillatory Dynamics in Alzheimer’s Disease: A Laminar Whole-Brain Model of Serotonergic Psychedelic Effects

**DOI:** 10.1101/2024.12.15.628565

**Authors:** Jan C. Gendra, Edmundo Lopez-Sola, Francesca Castaldo, Èlia Lleal-Custey, Roser Sanchez-Todo, Jakub Vohryzek, Ricardo Salvador, Ralph G. Andrzejak, Giulio Ruffini, the Alzheimer’s Disease Neuroimaging Initiative

## Abstract

Classical serotonergic psychedelics show promise in addressing neurodegenerative disorders such as Alzheimer’s disease by modulating pathological brain dynamics. However, the precise neurobiological mechanisms underlying their effects remain elusive. This study introduces a personalized whole-brain model built upon a laminar neural mass framework to elucidate these effects. Using multimodal neuroimaging data from thirty subjects diagnosed with mild to moderate Alzheimer’s disease, we simulate the impact of serotonin 2A receptor activation, characteristic of psychedelics, on cortical dynamics. By modulating the excitability of layer 5 pyramidal neurons, our models reproduce hallmark changes in EEG power spectra observed under psychedelics, including alpha power suppression and gamma power enhancement. These spectral shifts are shown to correlate strongly with the regional distribution of serotonin 2A receptors. Furthermore, simulated EEG reveals increased complexity and entropy, suggesting restored network function. These findings underscore the potential of serotonergic psychedelics to reestablish healthy oscillatory dynamics in the prodromal and early phases of Alzheimer’s disease and offer mechanistic insights into their potential therapeutic effects in neurodegenerative disorders.

## 1 Introduction

In the last couple of decades, classical serotonergic psychedelics (hereby, *psychedelics*) have re-emerged as potential treatments for a range of different neuropsychiatric disorders, yielding promising results [Nutt et al., 2020,Husain et al., 2023,Cameron and Patel, 2023]. Emerging evidence also points to the potential of psychedelics in treating neurodegenerative conditions, such as Alzheimer’s disease (AD) and related dementias [Winkelman et al., 2023, Garcia-Romeu et al., 2022, Vann Jones and O’Kelly, 2020, Kozlowska et al., 2022]. While current evidence is primarily preclinical, there is growing interest in investigating psychedelics as potential treatments for AD, with both high-dose and micro-dosing regimens being considered [Vann Jones and O’Kelly, 2020, Garcia-Romeu et al., 2022].

Serotonergic psychedelics primarily act as agonists of the serotonin 2A receptor (5-HT2AR) [Nutt et al., 2020, Cameron and Patel, 2023, Carhart-Harris and Friston, 2019, Carhart-Harris and Chandaria, 2023], though other receptors, such as 5-HT1AR, may also contribute to these effects [Warren et al., 2024, Hansen et al., 2022]. The intensity of these effects is linked to the degree of occupancy of these receptors [Madsen et al., 2019]. Among the different serotonin receptor (5-HT) subtypes, 5-HT2ARs are the most abundant in the cortex [Luppi et al., 2024] and show some regional heterogeneity [Nutt et al., 2020, Berger et al., 2009, Andrade, 2011]. 5-HT2A receptors modulate the excitatory effects of glutamate, particularly by enhancing the excitability of deep-layer pyramidal neurons to glutamate inputs (i.e., making them more susceptible to excitatory inputs typically associated with glutamate receptors) [Andrade, 2011]. This enhanced excitation dysregulates the spontaneous activity in cortical populations, increasing the entropy or complexity of ongoing brain activity and producing alterations in its power spectrum [Nutt et al., 2020, Carhart-Harris and Friston, 2019]. In particular, the power changes associated with 5-HT2AR activation disrupt the brain’s normal alpha rhythm dynamics, resulting in a widespread decrease in the mean PSD of the alpha band in the cortex [Nutt et al., 2020, Carhart-Harris and Friston, 2019, Carhart-Harris and Leech, 2014, Ort et al., 2023, Timmermann et al., 2023] and an increase in the cortical gamma band power [Timmermann et al., 2023, Smausz et al., 2022].

These mechanisms may be particularly valuable in the context of Alzheimer’s disease, where disrupted oscillatory patterns, reduced complexity, and impaired connectivity are hallmarks of the disorder. Specifically, these disruptions evolve across the stages of the disease, reflecting progressive impairments in neural dynamics. During the preclinical phase, alpha activity is disrupted [Nakamura et al., 2018,Gallego-Rudolf et al., 2024], while gamma power shows variability, with reports of increases, stability, or reductions [Gaubert et al., 2019, Rochart et al., 2020]. As the disease progresses into the prodromal phase, alpha power remains elevated [Nakamura et al., 2018]. During this phase, gamma power is thought to decline due to the dysfunction of parvalbumin (PV) interneurons, which are essential for generating gamma oscillations and maintaining network balance [Murty et al., 2021,Palop and Mucke, 2016]. In the mild to moderate stages, alpha power declines, gamma power becomes markedly reduced, and neural complexity decreases, reflecting growing impairments in synaptic function, network flexibility, and cognitive processing [Garcés et al., 2013, Abásolo et al., 2006, Sun et al., 2020]. For a more detailed discussion of AD physiopathology and oscillatory deficits, see Supplementary Material S1.

Psychedelics, acting through the mechanisms described above, could counteract these deficits by reducing alpha power, thereby alleviating hyper-synchronization and enhancing network flexibility. Their ability to increase gamma power is particularly relevant for addressing PV interneuron dysfunction, potentially restoring gamma oscillations and rebalancing excitatory-inhibitory networks [Palop and Mucke, 2016, Halberstadt, 2015]. Additionally, psychedelics enhance neural complexity, increasing entropy and promoting a more flexible and integrated network state [Carhart-Harris, 2018, Nutt et al., 2020, Ruffini et al., 2023], which may help counteract the rigidity and loss of complexity observed in Alzheimer’s disease. These effects suggest that psychedelics could be especially effective in the prodromal and mild stages of the disease, where compensatory mechanisms remain active, and neural circuits are less extensively damaged. A summary of how psychedelics may compensate for these deficits in AD is presented in Table 1.

**Table 1:** Compensation of AD deficits by serotonergic psychedelics, which suggest potential therapeutic value in early AD.

| Feature | AD Deficit | Compensation by psychedelics |
| --- | --- | --- |
| **Alpha Power** | Increased alpha power and alpha hypersynchronization (preclinical and prodromal) [Nakamura et al., 2018, López et al., 2014, Gallego-Rudolf et al., 2024]; eventual decrease in alpha peak frequency and alpha power in later stages [Passero et al., 1995, Benwell et al., 2020, Wang et al., 2015]. | Psychedelics reduce alpha power, increasing network flexibility and interconnectivity [Nutt et al., 2020, Carhart-Harris and Friston, 2019, Kometer et al., 2013]. Potentially valuable in early AD. |
| **Gamma Power** | Possible preclinical increases in gamma [Gaubert et al., 2019], decreased gamma activity in prodromal and later stages [Murty et al., 2021], impairing cognitive processes and memory [Palop and Mucke, 2016]. | Psychedelics increase gamma power, promoting oscillations associated with cognitive processing [Timmermann et al., 2023, Kometer et al., 2013, Smausz et al., 2022]. Potentially valuable after preclinical phase. |
| **Delta and Theta Power** | Increased power in delta and theta bands; hypersynchrony in slow oscillations [Nakamura et al., 2018, Passero et al., 1995, Chetty et al., 2024]. | Psychedelics reduce delta and theta power, countering hypersynchrony and restoring balance across frequencies [Pallavicini et al., 2019]. |
| **Complexity** | Decreased entropy and complexity of brain activity [Abásolo et al., 2006]. | Psychedelics increase entropy and brain signal complexity [Carhart-Harris, 2018, Carhart-Harris and Friston, 2019, Nutt et al., 2020, Ruffini et al., 2023]. |
| **Inhibitory-Excitatory Balance** | Impaired PV interneuron function disrupts gamma oscillations and leads to network instability [Palop and Mucke, 2016]. | Psychedelics engage 5-HT<sub>2A</sub>R on pyramidal neurons, indirectly improving PV interneuron function and rebalancing excitatory-inhibitory networks [Halberstadt, 2015]. |
| **Network Features** | Decreased global connectivity and increased modularity; network decoupling [Babiloni et al., 2020]. | Psychedelics increase global functional connectivity and reduce modularity, promoting integration and flexibility [Tagliazucchi et al., 2016]. |

A deeper understanding of the mechanisms underlying the effects of psychedelics is essential to develop such effective treatments [Carhart-Harris and Friston, 2019, Nutt et al., 2020,Cameron and Patel, 2023]. Psychedelics affect brain dynamics across scales, from sub- cellular mechanisms to whole-brain networks, making computational models valuable tools for exploring their therapeutic potential. Such mechanistic models have recently proven helpful in advancing our understanding of the mechanism of action of psychedelics [Deco and Cruzat, 2018, Kringelbach and Cruzat, 2020, Vohryzek et al., 2024, Vohryzek et al., 2022, Herzog et al., 2023, Mindlin and Herzog, 2023, Jobst et al., 2021, Luppi et al., 2022]. However, existing computational models primarily focus on macroscale brain dynamics, overlooking critical mesoscale and microscale dynamics, such as neuronal population interactions that mediate oscillatory changes. Moreover, there is a notable lack of computational studies examining the effects of psychedelics in the context of AD.

Mesoscale modeling may be well suited to reproduce such oscillatory alterations by psychedelics in AD, as it represents an intermediate level between the macro- and micro-scale and focuses on neuronal populations and local circuits within brain regions. Computational approaches such as neural mass models (NMMs) and neural field models are frequently used in mesoscale modeling to simulate the collective behavior of neuronal populations [Jansen and Rit, 1995, Cook et al., 2022]. The laminar NMM (LaNMM) framework is a natural extension of the neural mass formalism to produce coupled fast (gamma) and slow (alpha) oscillatory activity and simulate realistic electrophysiological signals [Ruffini et al., 2020, Sanchez-Todo and Bastos, 2023]. It has been recently applied in the context of AD [Sanchez-Todo et al., 2024], showing how disrupted PV-to-pyramidal connectivity can explain the PSD alterations observed in the process of neurodegeneration in AD. Additionally, we have used the LaNMM to investigate the role of cross-frequency coupling (CFC) in hierarchical predictive coding, showing how error evaluation and precision mechanisms deteriorate in AD due to disruptions of inhibitory interneuron function that impair the computation and gating of prediction errors [Ruffini et al., 2025]. By capturing multiband oscillatory dynamics, the LaNMM provides a unique framework for modeling neuronal processes in conditions like AD or under exposure to psychedelics, where disruptions in slow and fast oscillatory activity are prominent.

In this study, we investigate the effects of psychedelics at the whole-brain level and their therapeutic potential in AD using the LaNMM. With multimodal data from a cohort of AD subjects, we develop personalized hybrid brain models based on the LaNMM. We hypothesize that these whole-brain models can reproduce the spectral changes observed in empirical studies of psychedelic administration, specifically a pronounced decrease in alpha power and an increase in gamma power. Given the known disruptions in oscillatory activity in AD, we hypothesize that psychedelics may counteract key spectral abnormalities associated with the disease—particularly the imbalance between alpha and gamma oscillations. Furthermore, we predict that these changes will correlate with the spatial distribution of 5-HT2A receptor expression, supporting the notion that psychedelics modulate cortical excitability in a regionally specific manner. Finally, we explore whether psychedelics enhance neural complexity and entropy, which have been proposed as markers of cognitive flexibility and may contribute to their therapeutic effects.

## 2 Methods

An overview of the methodology followed in this study is shown in Figure 1. We first generated subject-specific whole-brain models based on diffusion magnetic resonance imaging (dMRI) and functional MRI (fMRI) data from AD subjects (Stage I). We then integrated the effects of psychedelics following a positron emission tomography (PET)-informed distribution of 5-HT2AR densities and study the effects of psychedelics at the whole-brain level in the cohort of AD subjects (Stage II). We used such models to evaluate the changes in oscillatory activity at the cortical level. Finally, we combined such whole-brain models with a biophysical layer to simulate realistic electrophysiological activity (i.e., EEG) under psychedelics (Stage III), and evaluate changes in alpha and gamma power and complexity metrics.

**Figure 1:**
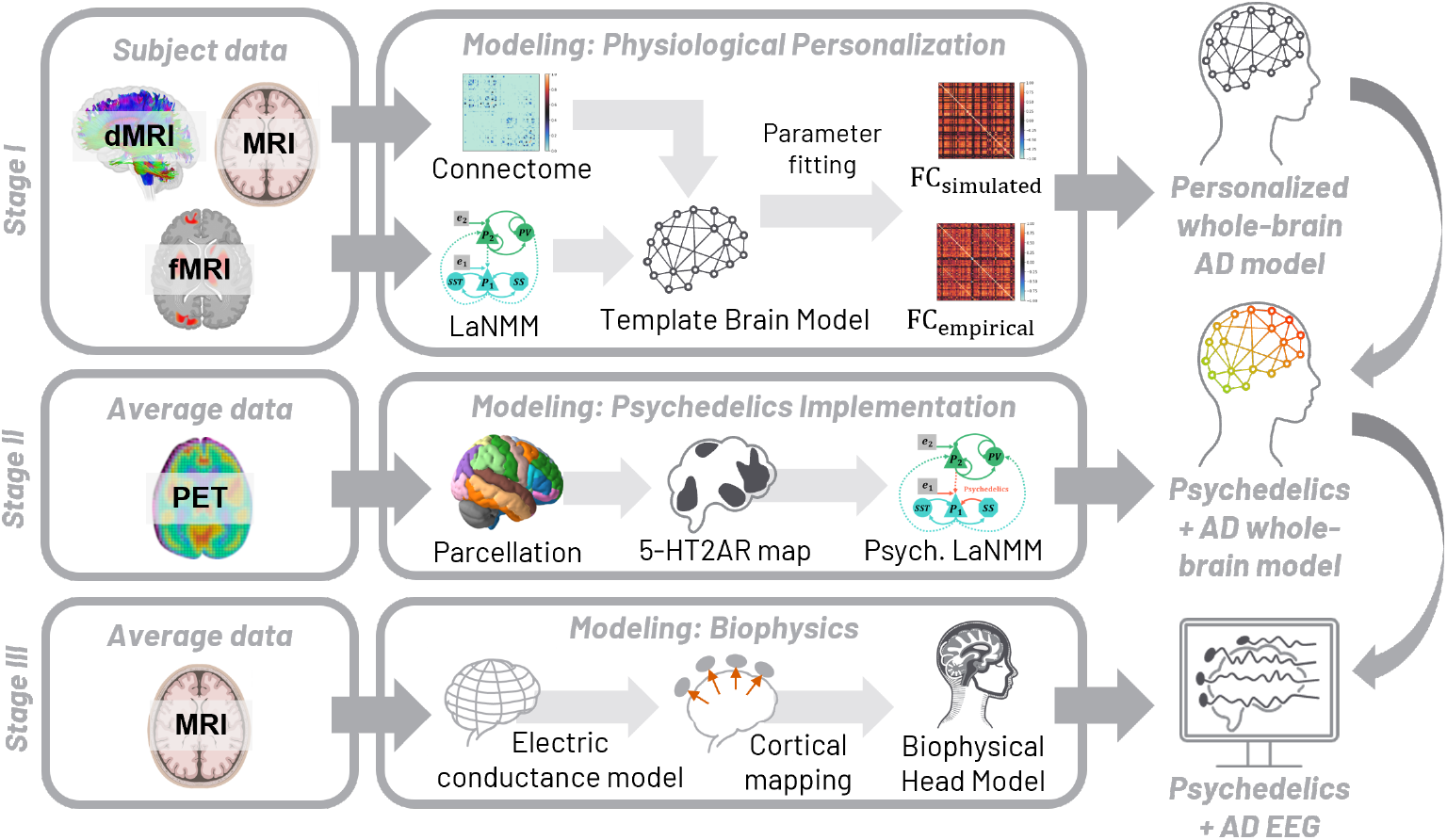
Overview of the methodology. — Pipeline for personalization of hybrid brain models of AD and the effects of psychedelics using structural MRI, dMRI, fMRI, and PET data. In Stage I, we generate personalized whole-brain models based on the LaNMM using multimodal data from a cohort of AD patients. In Stage II, we simulated the effects of psychedelics in the personalized models using the distribution of 5-HT2A receptors obtained from PET, and evaluated the oscillatory changes at the cortical level. In Stage III, we combined the whole-brain models with a biophysical head model to study the effects of psychedelics at the EEG level, including oscillatory changes and variations in complexity.

### 2.1 Laminar neural mass model implementation

The LaNMM consists of two coupled neural masses—a Jansen-Rit (JR) [Jansen and Rit, 1995] and a pyramidal-interneuron-gamma (PING) model [Börgers et al., 2008]— to simulate both slow and fast oscillations in a cortical column [Sanchez-Todo and Bastos, 2023]. The JR NMM model can generate slow oscillations in the alpha band (8–12 Hz) and is adjusted to oscillate at 10 Hz at baseline conditions. A variation of the PING model modified to oscillate at 40 Hz at baseline conditions produces fast oscillations in the gamma band (30-70 Hz).

The JR sub-circuit (Figure 2) consists of a population of pyramidal neurons (representing layer 5 pyramidal cells; *P*_1_) coupled via excitatory connections to a population of excitatory neurons (representing mostly spiny stellate neurons; *SS*) and to a population of slow inhibitory interneurons (representing somatostatin-expressing cells; *SST*). Both *SS* and *SST* populations are reciprocally coupled to *P*_1_ through excitatory and inhibitory synapses, respectively. On the other hand, the PING sub-circuit consists of a population of pyramidal cells (representing layer 2/3 pyramidal cells; *P*_2_) and a population of fast inhibitory interneurons (representing parvalbumin-positive cells; *PV*). External perturbations *e*_1_ and *e*_2_ are applied to *P*_1_ and *P*_2_, respectively, to represent the influence of other brain areas. The PING and JR sub-circuits are reciprocally coupled through excitatory connections between the pyramidal populations *P*_1_ and *P*_2_ and through the PING’s inhibitory population, thus constituting a single NMM. A schematic diagram can be found in Figure 2. The equations and model parameters for the LaNMM have been presented in previous work [Sanchez-Todo and Bastos, 2023] and are detailed in Supplement S2 and Table S1.

**Figure 2:**
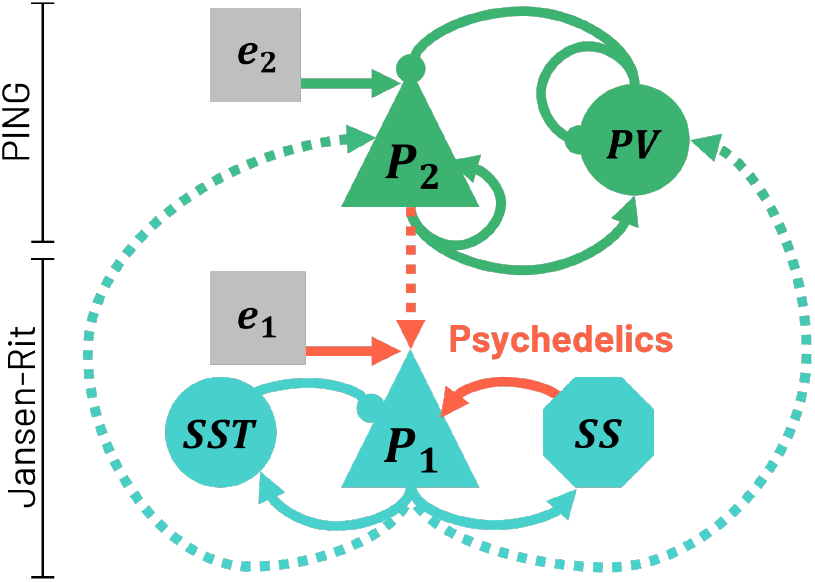
Schematic diagram of the LaNMM depicting its neuronal populations and the connections between them. Red arrows indicate the synapses affected by 5-HT2AR activation due to psychedelics in the model. Dashed lines represent connections between the PING and Jansen-Rit parts sub-circuits.

#### 2.1.1 Implementation of psychedelics effects

The increased excitability to glutamate inputs observed under psychedelics can be implemented in the model as an increase of the average synaptic gain *A*_*L*5*P*_ parameter of excitatory connections to L5 pyramidal populations in the LaNMM. This parameter determines the extent to which the average firing rate of a population of neurons increases or decreases in response to an input membrane potential perturbation. Three glutamatergic connections to L5 pyramidal neurons are present in the LaNMM: connections from the *SS* and *P*_2_ populations to *P*_1_, and external noise perturbation *e*_1_ to *P*_1_. Hence, we model 5-HT2AR activation by transforming receptor stimulation into changes of the excitatory synaptic gain *A*_*L*5*P*_ in those connections.

We studied the dynamics of a single uncoupled LaNMM for different *A*_*L*5*P*_ values in the three targeted connections. For simplicity, we modified this parameter equally for all these synapses, assuming that the activation of 5-HT2ARs affects similarly all L5 pyramidal synapses in a particular cortical column. We have used the average membrane potential activity of the pyramidal populations to study the dynamics of the uncoupled LaNMM, given that pyramidal neurons are thought to be the main contributors to the potential recorded in electrophysiological signals such as EEG [Nunez and Srinivasan, 2006, Buzsáki et al., 2012].

The model was simulated for each gain value *A*_*L*5*P*_ ∈ [0, 10] mV in increasing steps of 0.05 mV. In each simulation, *A*_*L*5*P*_ was simultaneously and equally increased in all three target connections, and the LaNMM was run for a duration of 40 s with a sampling frequency *f*_*s*_ = 1000 Hz. Then, the PSD of the membrane potential of each pyramidal population *P*_1_ and *P*_2_ was computed for the last 30 s of the simulation to allow the model’s convergence to the steady state. Throughout the study, the PSD is calculated using Welch’s method [Welch, 1967] with a segment length *L* = 1000 samples.

### 2.2 AD whole-brain models personalization

We created personalized whole-brain models of a cohort of 30 AD subjects using the LaNMM (Stage I in Figure 1).

#### Patient data

We used data from 30 AD subjects from the Alzheimer’s Disease Neuroimaging Initiative 3 (ADNI3) database (https://adni.loni.usc.edu/). The ADNI was a multicenter study initiated by Michael W. Weiner in 2004, which focused on the early diagnosis and follow-up of AD. We selected subjects diagnosed with AD for which the multimodal data required was available and with sufficient variability in age, gender, and cognitive function. The cohort selected consisted of 13 women and 17 men, with a mean age of 73.6 years (SD: 7.32). The average Mini-Mental State Examination (MMSE) score, a standard measure of cognitive function, ranged from 17 to 30 (mean: 22.5, SD: 2.7), reflecting variability in the extent of cognitive decline among the subjects selected. The data acquisition parameters for each modality used in this study and the patient cohort selected are provided in S3.

#### Data processing

Each modality underwent a dedicated processing pipeline to generate subject-specific inputs for the whole-brain models. T1-weighted MRI data was used to create a subject-specific Desikan-Killiany (DK-68) cortical atlas [Desikan et al., 2006]. The MRI processing pipeline, based on FreeSurfer [Fischl et al., 2004], began with cortical reconstruction using the *recon-all* function to extract the pial surface. The DK68 parcellation, provided by FreeSurfer, was mapped to the subject-specific brain structure. This volumetric image served as the basis for processing dMRI and rs-fMRI data.

Diffusion MRI data was processed to calculate the subject’s structural connectome (SC). Preprocessing steps included denoising, artifact correction (e.g., eddy currents), and estimating the fiber orientation density using constrained spherical deconvolution. Probabilistic tractography was then performed with MRtrix3 [Tournier et al., 2019], generating 20 million streamlines constrained to start and terminate in gray matter parcels defined by the MRI parcellation. The resulting structural connectome matrix quantified the number of streamlines connecting each pair of parcels and was normalized by the total number of fibers and the number of parcels to generate a connectivity matrix for the subject.

BOLD rs-fMRI data was preprocessed using fMRIPrep 22.0.0 [Esteban et al., 2019], which utilizes Nipype 1.8.3 [Gorgolewski et al., 2011]. For anatomical data, T1-weighted images were corrected for intensity non-uniformity (N4BiasFieldCorrection) and skull-stripped using ANTs [Avants et al., 2009]. Tissue segmentation was performed with FSL FAST, and brain surfaces were reconstructed with FreeSurfer. Functional data preprocessing included motion correction (FSL mcflirt), slice-timing correction (AFNI 3dTshift), co-registration to T1-weighted images (FreeSurfer bbregister), and normalization to MNI templates using ANTs. Confound regressors, including motion parameters, CompCor components, and framewise displacement, were calculated to facilitate nuisance regression. Detailed preprocessing workflows and parameters are described in the supplementary materials and in the fMRIPrep documentation (https://fmriprep.org).

#### Model fitting

In our framework, whole-brain models represent the human cortex as a network of coupled NMMs, with each node representing the average activity of a cortical parcel through a LaNMM and each link between them modeling the white matter fibers connecting these parcels. The connectome (obtained from dMRI data as described above) was used to provide the connectivity between nodes. *P*_1_ and *P*_2_ populations of different parcels *i, j* were connected as follows: 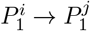 (lateral connection), 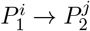 (feedback or descending connection) and 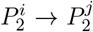 (feedforward or ascending connection) with relative weights of 0.5, 0.5 and 1 respectively. Further details about the long-range connectivity are given in Supplementary Material S2.2.

Whole-brain models were personalized to match each subject’s empirical functional connectivity (FC) obtained from their BOLD rs-fMRI, ultimately tailoring the models to reflect individual neural dynamics. Synthetic BOLD data from the whole-brain models was generated using the Balloon-Windkessel model [Friston et al., 2000], and FC matrices were computed using Pearson’s correlation coefficient (PCC) between the BOLD signals. Each whole-brain model was simulated for five different noise realizations for a duration of 300 s with a sampling frequency of *f*_*s*_ = 1000 Hz. The first 5 seconds of the simulation were discarded to account for transient dynamics and ensure the models reached a steady state. The fourth-order Runge-Kutta method [Press and Teukolsky, 1992] (RK4) was used to solve the model’s differential equations.

Model optimization involved tuning the global coupling gain *G* (which scales the connectome) and the noise received by the pyramidal populations of each region (in particular, the ratio between common and homotopic noise standard deviations (SDs) *σ*_*c*_*/σ*_*i*_, see Supplementary Material S2.3). A grid search was performed to select the optimal values of *G* and noise that minimized the root mean squared error (RMSE) between simulated and empirical FC matrices for each subject. The best value across noise realization was selected. An additional constraint was applied, requiring the PCC between simulated and empirical FC matrices to be higher than the PCC between the empirical FC and the structural connectivity matrix. This ensured that the models captured FC patterns beyond those explained by anatomical connectivity.

#### 2.3 Whole-brain models under psychedelics

The 30 subject-specific whole-brain models described in Section 2.2 were adapted to reproduce the activation of 5-HT2A receptors (Stage II in Figure 1). Following the approach described in Section 2.1, we implemented the activation of 5-HT2A receptors as changes in the synaptic gain of all excitatory inputs to *P*_1_ pyramidal population (*A*_*L*5*P*_) in every LaNMM of the brain network. This includes connections from the *SS* and *P*_2_ populations to *P*_1_, the common and homotopic external inputs (see Supplementary Material S2.3), and all *P*_1_ → *P*_1_ long-range cortico-cortical inputs.

To modulate the average synaptic gain of the connections involved in the psychedelic effect, we used 5-HT2A receptor density data informed by the high-resolution *in vivo* atlas of the human brain serotonin system measured with PET in Beliveau et al. [Beliveau et al., 2017]. We refer the reader to that paper for further details on the PET and MRI acquisition parameters and the participant’s demographics. We parcellated the 5-HT2AR average density map into the DK-68 cortical atlas as detailed in Supplementary Material S4. We then normalized the density values *B*_*max*_ to fall between 0 and 1. This provides us with the relative spatial distribution of 5-HT2A receptors between brain areas, defining the amount of serotonergic stimulation each brain area experiences under psychedelic drugs.

To translate the receptor density map into average synaptic amplitude gain values of glutamatergic connections to *P*_1_ (*A*_*L*5*P*_), we define the neuromodulation relation

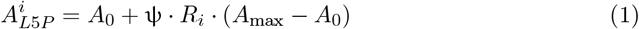

where ψ is the psychedelic dose (0 at baseline, 1 under psychedelics) and *R*_*i*_ ∈ [0, 1] is the normalized parcel’s receptor density. *A*_0_ = 3.25 mV is the nominal gain, i.e., the average synaptic amplitude (PSP) gain of pyramidal neurons at baseline conditions [Jansen and Rit, 1995]. *A*_max_ = 3.75 mV is the maximum gain, i.e., the gain at which the PSD decay of the alpha band in the *P*_1_ population ceases to be linear when increasing *A*_*L*5*P*_ in an uncoupled LaNMM, determined as explained in Section 3.1. Hence, this maps the receptor densities between *A*_0_ and *A*_max_ and maintains a linear relationship between *R* and *A*_*L*5*P*_ . The normalization applied to *B*_*max*_ assumes that the minimum density of receptors corresponds to the nominal gain *A*_0_. Of note, we use the same *A*_max_ value in all parcels and apply the same spatial distribution of 5-HT2AR densities to each subject.

Each subject’s whole-brain model of psychedelic effects was simulated for a duration of 40 s with a sampling frequency *f*_*s*_ = 1000 Hz. The RK4 method was used to solve the model’s differential equations. The first 10 seconds of the simulation were disregarded in all subsequent analyses to account for the transient time of the models and their convergence to the steady state. We computed the parcel-averaged power spectrum of each subject’s membrane potential and conducted Wilcoxon signed-rank tests in each brain band (delta: 0.5–4 Hz, theta: 4–8 Hz, alpha: 8–12 Hz, beta: 12–30 Hz, and gamma: 30-70 Hz), comparing the mean PSD in each frequency band across subjects between the baseline and psychedelic conditions. We used the Wilcoxon signed-rank test as a non-parametric alternative to the paired t-test, given that the data did not conform to a normal distribution according to the Kolmogorov–Smirnov test (Bonferroni-corrected p<0.05 in all frequency bands and conditions).

To assess whether the oscillatory changes correlated with the distribution of 5-HT2A receptors obtained from PET, we measured the PCC between the density of 5-HT2A receptors in each parcel and the absolute PSD difference (*PSD* _Post_−*PSD* _Pre_) in both the alpha and gamma bands in that parcel. Parcels with 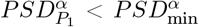 at baseline were excluded from the analysis since their alpha power was not expected to show relevant variations due to changes in receptor density.

### 2.4 Simulation of EEG recordings

We generated synthetic EEG data for each subject using *hybrid brain models*, which combine the physiological whole-brain models developed in the previous section with a biophysical head model template [Miranda et al., 2013,Ruffini et al., 2013,Ruffini et al., 2014] (Stage III in Figure 1). The process of EEG generation from the simulated activity in the whole-brain model NMMs can be divided into two steps: 1) generation of the equivalent dipole strength over time from NMM activity, 2) generation of EEG signals from the dipole strength.

To translate NMM activity into the equivalent dipole strength, we used the mesoscale physical model of volume conduction and the neuron compartment models of current generation from Mercadal et al. (2023) [Mercadal et al., 2023], which were developed based on the laminar framework developed in Sanchez-Todo et al. (2023) [Sanchez-Todo and Bastos, 2023]. In brief, a multi-compartmental modeling formalism and detailed geometrical models of pyramidal cells are used to simulate the transmembrane currents due to synaptic inputs into different cortical layers. These results are linearly combined to estimate the total current source densities (CSDs) generated due to the neural activity in pyramidal populations, which are considered the main current generators [Nunez and Srinivasan, 2006, Buzsáki et al., 2012]. The CSDs are then used to calculate the equivalent dipole strength over time associated with the neural activity in each parcel of the subject-specific whole-brain model.

To generate scalp EEG signals from the dipole activity, we combined the personalized whole-brain model with a template biophysical head model, creating a hybrid brain model [Sanchez-Todo et al., 2018]. Supplementary Material S5 details the process of template head model creation. We used such a head model of electrical conduction to compute cortical mappers for EEG and generate the EEG signals in each electrode from the equivalent dipole strength computed in the personalized whole-brain models.

We computed the power spectrum of each EEG channel averaged across subjects. We then conducted Wilcoxon signed-rank tests in the alpha and gamma bands, comparing the mean band PSD of both conditions. The use of the Wilcoxon signed-rank test was again justified by the non-normal distribution of the data, confirmed by the Kolmogorov-Smirnov test (Bonferroni-corrected p<0.05 in all EEG channels and conditions).

#### 2.4.1 Complexity and entropy of simulated EEG

To capture the complexity of the simulated EEG signals obtained from the personalized whole-brain models under psychedelics, we used practical approximations of algorithmic complexity based on Lempel–Ziv compression and entropy rate, as detailed in Ruffini et al. (2019) [Ruffini, 2017, Ruffini et al., 2019].

We estimated the *Lempel-Ziv complexity* of the EEG data signals using the Lempel-Ziv-Welch (LZW) method [Cover, 1999,Ruffini, 2017,Ruffini et al., 2019,Welch, 1984]. We used a single 5-second epoch of simulated broadband EEG data (0.5–100 Hz) generated from each subject’s personalized computational model for this calculation and the subsequent entropy rate analysis, as the intrasubject variability of the synthetic data was negligible. The 5-second epoch of each channel was binarized based on the median value of the epoch: each value was converted into a ‘0’ if it was below the median and into a ‘1’ otherwise [Ruffini, 2017]. The binarized data of all channels was concatenated and the LZW method was applied to the resulting string. A natural way to normalize the description length resulting from the LZW method, *l*_*LZW*_, is to divide it by the original string length *n, ρ*_0_ = *l*_*LZW*_ */n*, with units of bits per character. We computed *ρ*_0_ following this method for each subject.

To complement this analysis, we computed the related *entropy rate* of the binarized EEG signals for each subject. For a stationary stochastic process {*X*_*i*_ }, the entropy rate ℋ (*X*) is defined as

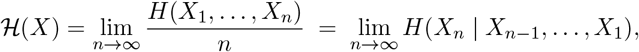

where *H*_*n*_(*X*) = *H*(*X*_1_, …, *X*_*n*_) is the joint entropy and *H*(*X*) = −𝔼_*X*_[log *P* (*X*)]. For a Markov process of order *n*, the entropy rate simplifies to the conditional entropy given the previous *n* states. In the special case of *n* = 0 (an independent and identically distributed process), the entropy rate reduces to the entropy of a single symbol, *H*(*X*_*i*_). Following these definitions, we computed the entropy rate for Markov orders *n* = 0, …, 5. [Cover, 1999, Ruffini et al., 2019]

Finally, we calculated the *spectral entropy* of the 5-second epoch of simulated EEG as an alternative measure of neural entropy under baseline and psychedelic conditions. To achieve this, for each subject, the PSD was averaged across channels and normalized to sum to 1, thereby transforming it into a probability distribution function (PDF). The spectral entropy was then computed for each channel using Shannon’s entropy, *H*(*p*) = − ∑*p*_*i*_ · log *p*_*i*_, where *p*_*i*_ is the PSD normalized to unit sum.

## 3 Results

### 3.1 Uncoupled LaNMM spectral profile

We first studied the power spectrum of pyramidal populations *P*_1_ (generator of alpha oscillations) and *P*_2_ (generator of gamma oscillations) in an uncoupled LaNMM, and assessed the LaNMM’s ability to simulate the cortical spectral alterations resulting from 5-HT2AR psychedelics activation in this range.

Figure 3 shows the mean PSD in the alpha and gamma bands of populations *P*_1_ and *P*_2_ for different synaptic gains *A*_*L*5*P*_ ∈ [0, 10] mV. We can observe a window of synaptic gain values *A*_*L*5*P*_ ∈ [2.75, 4.15] mV for which both alpha and gamma oscillations are present in the LaNMM (as further detailed in Supplementary Material S6).

**Figure 3:**
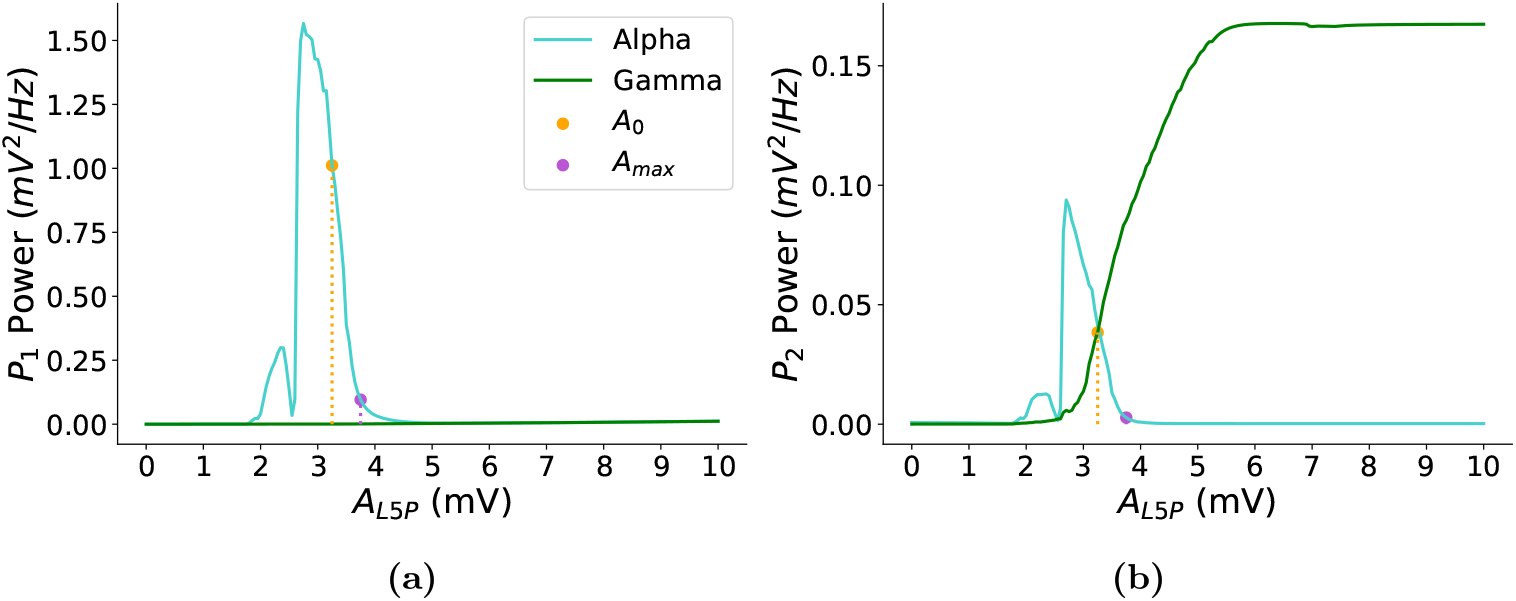
Uncoupled LaNMM spectral profile. Mean PSD of the membrane potential of *P*_1_ **(a)** and *P*_2_ **(b)** in the alpha band (blue) and gamma band (green) for different values of synaptic gain *A*_*L*5*P*_ . The vertical dashed lines indicate the nominal gain *A*_0_ (orange) and the maximum gain *A*_max_ (purple), respectively.

The results show a main peak in the mean PSD of *P*_1_’s alpha band coinciding with the start of the gamma oscillations in *P*_2_ at *A*_*L*5*P*_ = 2.75 mV. The decay of alpha activity in *P*_1_ starts concurrently with the growth of gamma activity in *P*_2_, indicating a reciprocal and inverse relationship to synaptic gain changes between the two pyramidal populations for *A*_*L*5*P*_ ∈ [2.75, 4.15] mV. The nominal synaptic gain *A*_0_ = 3.25 mV used in Sanchez-Todo et al. (2023) [Sanchez-Todo and Bastos, 2023] and Jansen and Rit (1995) [Jansen and Rit, 1995] falls within this interval. This indicates that at baseline conditions (no psychedelics applied), *P*_1_ oscillates in the alpha band and *P*_2_ in the gamma band. The decay of alpha power in *P*_1_ becomes nonlinear at around 3.75 mV, with a PSD in alpha of about 0.1 mV^2^/Hz. We will use this value as the maximum gain of the pyramidal populations in our model, *A*_max_ = 3.75 mV (corresponding to a minimum PSD in the alpha band of 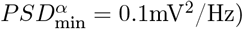.

Therefore, the decrease in alpha power in *P*_1_ and the increase in gamma power in *P*_2_ can be reproduced in the model through the selective increase of glutamatergic gain of synapses targeting L5 pyramidal neurons, associated with higher 5-HT2AR densities. The inverse relationship between alpha and gamma power is sustained within our regime of interest *A*_*L*5*P*_ ∈ [*A*_0_, *A*_max_]. It is worth mentioning that the most significant changes induced by psychedelics in the PSD magnitude in our LaNMM are specific to the alpha and gamma bands within the gain range under study in both *P*_1_ and *P*_2_ populations (Figure S6).

### 3.2 Whole-brain model personalization

The personalization of whole-brain models for the 30 AD subjects resulted in high similarity between the subjects’ rs-fMRI BOLD FC and the synthetic FC obtained with the fitted model, with an average PCC across subjects of 0.45 and an average RMSE of 0.18 after model personalization.

For all subjects, we found higher similarity (captured with RMSE and PCC) of the empirical FC with the model-generated FC than with the structural connectivity, confirming that the models were able to explain the functional data beyond the information provided by the subject-specific connectome alone (Figure 4a). The values of PCC, RMSE, and fitted parameters for each subject can be found in Supplementary S7, as well as the empirical and synthetic FC matrices for each subject.

**Figure 4:**
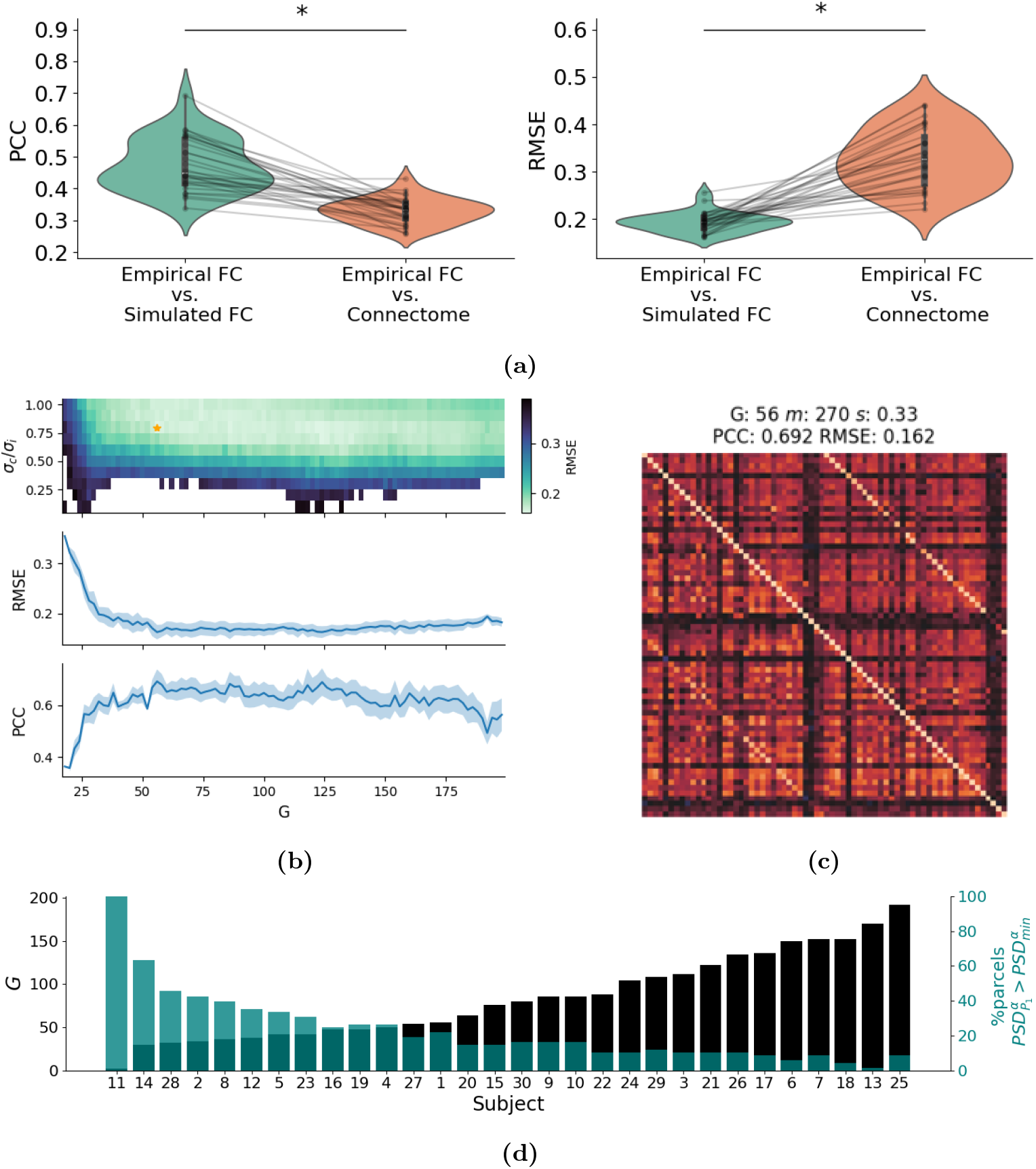
Whole-brain model fitting results. **(a)** PCC and RMSE between the empirical FC and simulated FC compared with the PCC and RMSE of the empirical FC and the structural connectome. Asterisks denote statistical significance in paired t-tests. **(b)** Example of model personalization (subject 1): RMSE between empirical and synthetic FC for different values of the parameters *G* and *σ*_*c*_*/σ*_*i*_ (top), RMSE and PCC as a function of the G value (center and bottom). **(c)** For the same subject, comparison between the empirical rs-fMRI BOLD FC (lower triangular matrix) and the synthetic FC obtained with the fitted model (upper triangular matrix). **(d)** Optimal *G* parameter found for each subject-specific model (from lowest to highest) and number of DK-68 parcels of each model with meaningful alpha activity in 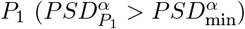.

As an example of model parameter optimization, Figure 4b shows the grid search performed to find the minimum value of RMSE for different combinations of *G* and *σ*_*c*_*/σ*_*i*_ for one subject model personalization. The FC matrices obtained from the synthetic BOLD signal for the optimal parameter selection for this subject are shown in Figure 4c and compared with the subject’s BOLD FC, showing the similarity between both matrices.

We then analyzed the optimal *G* parameter values found for each subject and their relationship to the alpha activity at baseline, which is crucial for the simulation of the effects of psychedelics in the model. Figure 4d shows the *G* parameter across subject models and the percentage of parcels in each model that are above the threshold of mean PSD for *P*_1_’s alpha band 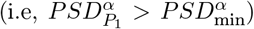. We ordered the values according to the magnitude of the personalized *G* parameter to facilitate the visualization. The high negative correlation observed between alpha activity and *G* (*r*_*P CC*_ = −0.80, *p <* 0.0001) evidences that personalized models with higher *G* values have less overall alpha activity in the brain. Since *G* controls the global excitation of the brain model, the higher *G* is, the more the *P*_1_ population activity approaches the region where alpha activity is lost (see Figure S5).

Of note, subject 11 showed an abnormally low *G*, which resulted in alpha oscillations in all cortical areas due to a globally diminished excitability. Upon deeper inspection, we noted that for this subject the grid-search selected parameter values resulted in total disconnection between nodes, leading to non-realistic dynamics. On the other hand, subject 13 showed an abnormally low alpha activity at baseline with only one alpha-active parcel, which is strongly related to its high *G* value. Further analysis revealed that all subjects except for subject 13 were responsive to 5-HT2AR stimulation. For these reasons, we excluded both subjects 11 and 13 from the calculations and analysis in the next sections.

It is worth mentioning that, on average, only 21% of the parcels have meaningful alpha activity at baseline (that is, above 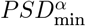) in the personalized models. Nevertheless, the models are able to reproduce significant changes in the mean alpha band activity following the simulation of 5-HT2AR activation, as shown in the next section.

### 3.3 Whole-brain spectral profile under psychedelics

Next, we studied the power spectrum of the AD whole-brain models and the spectral changes caused by psychedelics at the brain network level.

Figure 5 displays the power spectrum in *P*_1_ and *P*_2_ averaged across parcels and subjects and the results of the Wilcoxon signed-rank test. Consistent with our findings for the uncoupled LaNMM, clear peaks are observed in the alpha and gamma bands in the baseline condition. Moreover, the statistical analyses revealed a significant decrease in mean alpha power in both *P*_1_ (-61%) and *P*_2_ (-44%) and a significant increase in mean gamma power in both *P*_1_ (-25%) and *P*_2_ (-3.5%). The level of significance across tests was Bonferroni-corrected. The tendency towards a decrease of alpha power and an increase of gamma power in both pyramidal populations was consistently observed in all subjects, as shown in Figure 5b.

**Figure 5:**
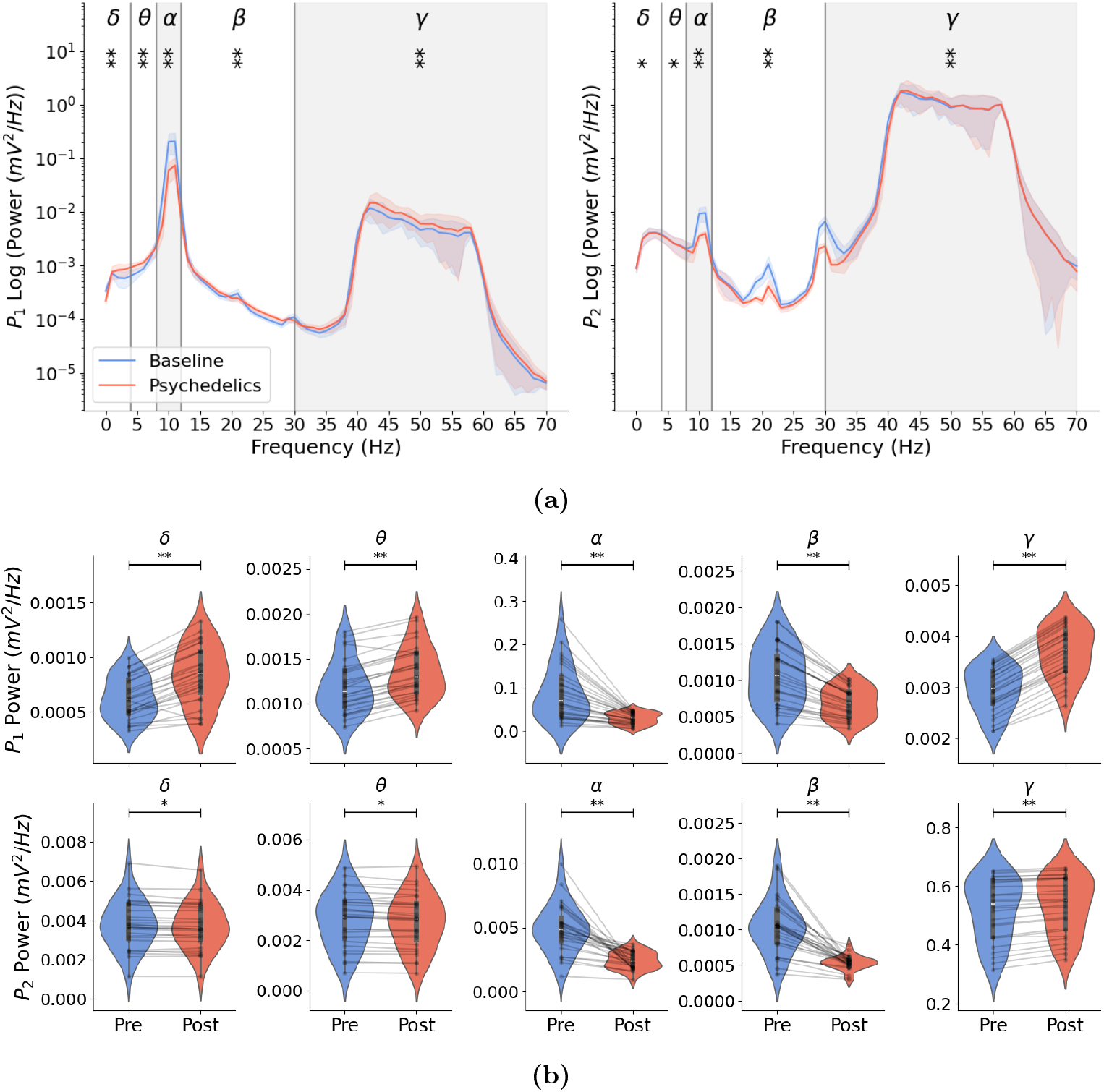
Whole-brain models reproduce the power spectrum alterations associated with psychedelics at the cortical level. **(a)** Power spectrum of *P*_1_ (left) and *P*_2_ (right) membrane potential averaged across brain parcels and subjects. Shaded areas represent the median absolute deviation across subjects, and solid line curves represent the average across subjects. The significance level (Bonferroni-corrected) of the average PSD difference between conditions in each band is denoted by the number of asterisks (*p < 0.05, **p < 0.0001). **(b)** Parcel-averaged membrane potential PSD of *P*_1_ (top) and *P*_2_ (bottom) in different bands across subjects. Black bars represent the interquartile range, and a central white dot indicates the median PSD. Individual dots connected by grey lines represent the mean power for each subject. Asterisks denote statistical significance as described above.

Although our model was specifically designed to produce alpha and gamma oscillations, we observed significant alterations across all bands and pyramidal populations under psychedelic conditions. Alpha and beta oscillations decreased on average with 5-HT2AR stimulation, while gamma oscillations increased. Nevertheless, the delta and theta bands showed discrepancies between the two pyramidal populations since their mean power consistently increased in *P*_1_ but decreased in *P*_2_ in every subject. The large standard deviation seen for the alpha band is consistent with the high variability seen in alpha power and *G* values between different subjects in Figure 4d.

Alterations in the beta band result from the harmonics of alpha oscillations (see Figure S6), while changes in the theta and delta frequency bands mainly result from the external noisy input introduced to population *P*_1_, a connection also susceptible to 5-HT2AR stimulation. The frequency power analysis also highlights that the greatest magnitude change in PSD is specific to the alpha and gamma frequency bands, which are the main focus of our study.

### 3.4 Influence of 5-HT2AR distribution on oscillatory activity

The changes in the spatial distribution of alpha and gamma activity across the brain are expected to relate to the applied PET-informed distribution of neurotransmitter densities. To evaluate this, we measured the PCC between the parcels’ density of 5-HT2A receptors and their absolute PSD difference (*PSD* _Post_ − *PSD* _Pre_) in both the alpha and gamma bands. Figure 6 shows the results of the analysis averaged across subjects (i.e., PSD difference of the parcel averaged across subjects). We found strong correlations for both the alpha band (*P*_1_ *r*_PCC_ = −0.38, *p <* 0.009) and the gamma band (*P*_1_ *r*_PCC_ = 0.91, *p <* 0.0001; *P*_2_ *r*_PCC_ = −0.49, *p <* 0.0001), except for the alpha power in *P*_2_ (*r*_PCC_ = − 0.04, *p* = 0.7). This suggests that regions with the highest expression of 5-HT2A receptors were most affected by psychedelics in the model.

**Figure 6:**
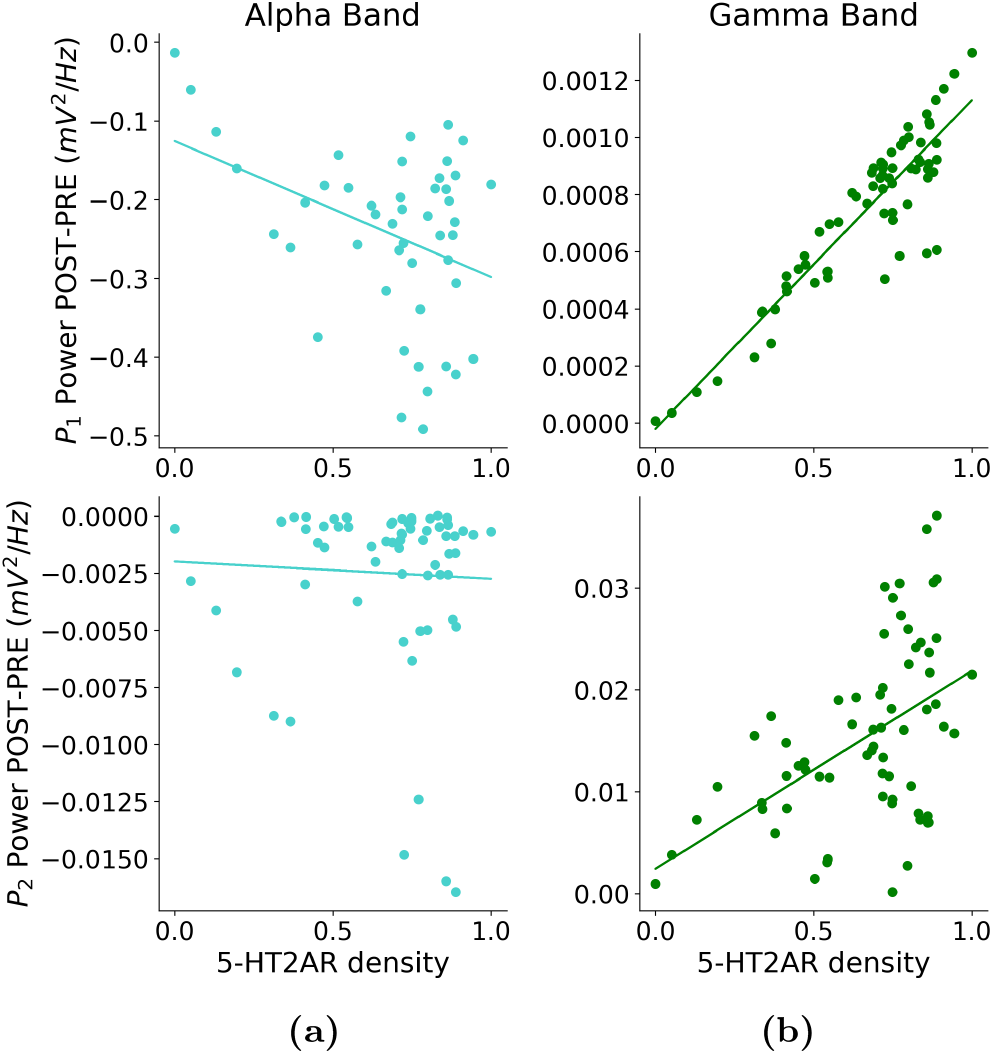
Spectral alterations under psychedelics reflect the spatial distribution and density of 5-HT2A receptors. Subject averaged parcel alpha band **(a)** and gamma band **(b)** membrane potential PSD difference (*PSD* _Post_ − *PSD* _Pre_) against parcel 5-HT2AR density in *P*_1_ (top) and *P*_2_ (bottom). Each dot represents a parcel, solid lines showing linear regression fits.

### 3.5 Model-generated EEG alpha and gamma power

Finally, we studied the power spectrum alterations induced by psychedelics in EEG data. Figure 7a shows the subject-averaged topographic map of the power in the alpha and gamma bands computed from the simulated EEG signals in the AD whole-brain models with and without psychedelics. The subject-averaged PSD difference between the baseline and psychedelic conditions is shown in Figure 7b, along with the statistical significance of the changes in each EEG electrode given by the analysis described above.

**Figure 7:**
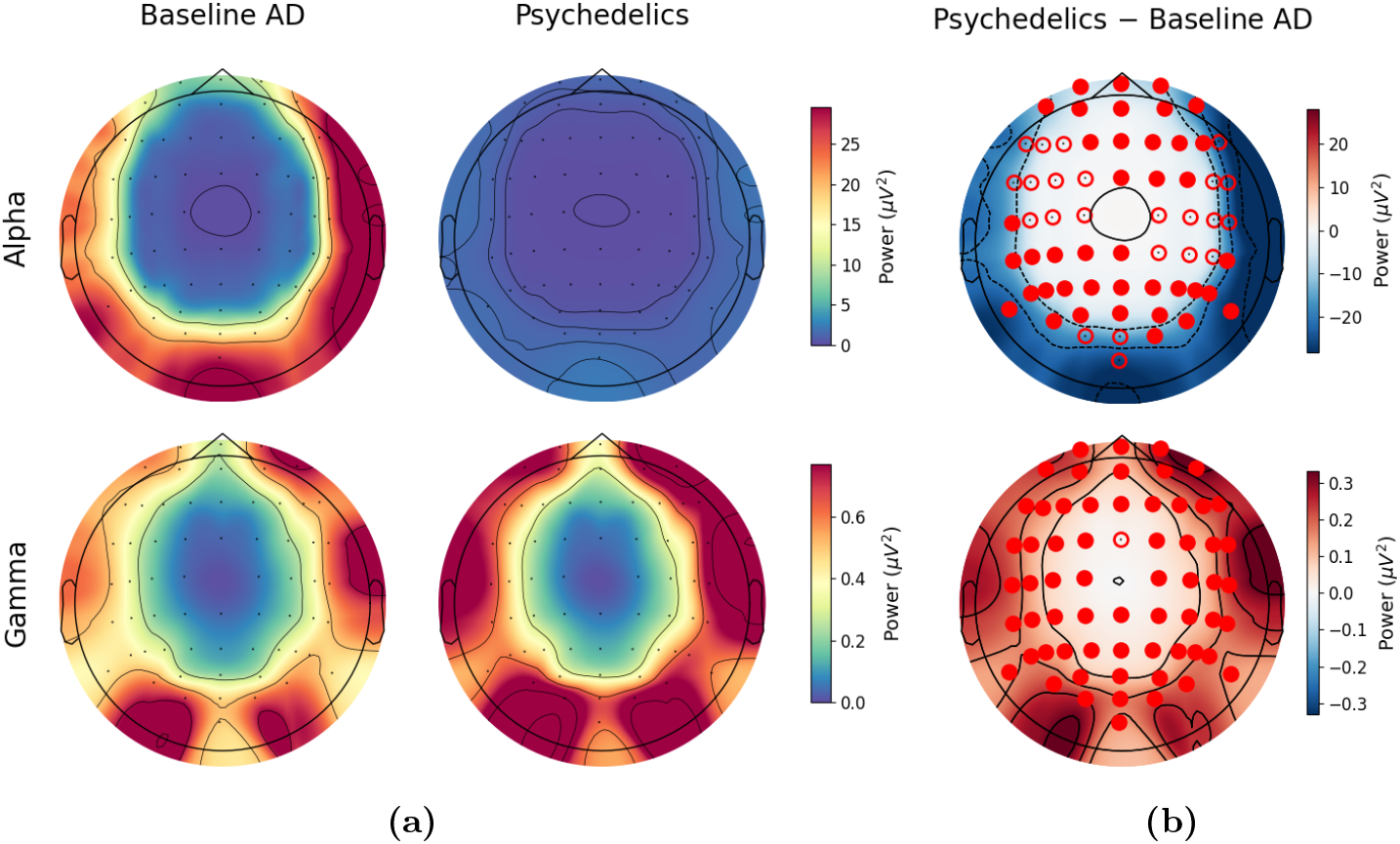
Model reproduces EEG alpha and gamma changes following 5- HT2AR stimulation with psychedelics. **(a)** Subject-averaged topographic map of the mean alpha and gamma band’s EEG PSD for the AD whole-brain models with and without stimulation by psychedelics. Black dots denote the recording location of the EEG electrodes. **(b)** Subject-averaged EEG PSD difference between the baseline AD models and the models with psychedelics effects (psychedelics − baseline). The statistical significance (Bonferroni-corrected) of the PSD changes in each EEG channel is denoted by an empty red dot (p < 0.05) or a filled red dot (p < 0.001) on top of the corresponding electrode.

Consistent with our previous results, the EEG shows a brain-wide decrease of alpha power in the spontaneous brain activity of all the modeled subjects (62% mean alpha PSD decrease with respect to baseline averaged over subjects and EEG channels). This decrease is significant across all EEG channels, and highly significant decreases in alpha power (p < 0.001) were consistently observed in all subjects and in all brain lobes. On the other hand, we found a significant increase in the EEG gamma band power after 5-HT2AR stimulation (37% mean gamma PSD increase with respect to baseline averaged over subjects and EEG channels). This increase is significant across all EEG channels and was consistently observed in all subjects.

It is worth mentioning that the cortical distribution of alpha and gamma activity in the synthetic EEG may not fully align with the spectral features typically observed in the EEG of healthy or AD individuals. The reduced oscillatory activity in both alpha and gamma frequencies in the parietal regions may be explained by the fact that the wholebrain models developed in this study were not personalized based on EEG data but on fMRI data. Inspection of the connectivity matrix revealed that this reduced EEG power can be mostly explained by a saturation of the connections to the parietal lobe. Indeed, the parietal lobe acts as a network hub, and the neural populations in this lobe receive high inputs and consequently often show saturated activity in the model. This hinders the emergence of oscillatory activity in alpha and gamma frequencies, resulting in the decreased alpha and gamma power observed in the parietal and parieto-frontal areas.

### 3.6 Simulated EEG complexity and entropy

Next, we used different metrics to capture the complexity of the simulated EEG signals obtained from the personalized hybrid brain models under psychedelics. Given the non-linear dynamics of EEG signals, we selected metrics that estimate upper bounds on algorithmic complexity (see Section 2.4.1). The normalized LZW complexity (*ρ*_0_) of EEG signals in subjects under psychedelics was significantly higher than at baseline (Figure 8a), demonstrating that the oscillatory changes induced by psychedelics in the whole-brain models lead to increased complexity of the signals.

**Figure 8:**
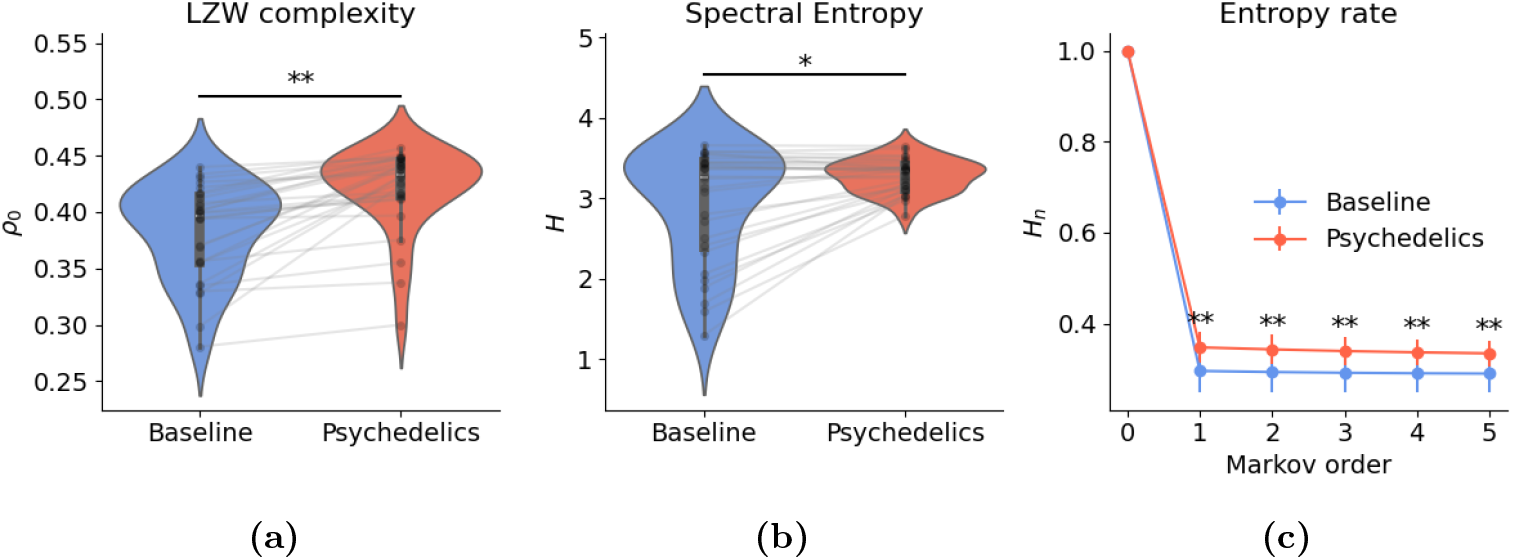
Model reproduces complexity and entropy changes under psychedelics. **(a)** Normalized LZW complexity of broadband EEG for the whole-brain models at baseline and under psychedelics. Black bars represent the interquartile range, and a central white dot indicates the median complexity. Individual dots connected by grey lines represent the signal complexity for each subject. **(b)** Spectral entropy of broad- band EEG signals for both conditions. **(c)** Entropy rate for different Markov orders of broadband EEG signals in the baseline and psychedelics conditions. Error bars indicate the standard deviation across subjects. In all cases, the significance level of the difference between conditions resulting from a paired t-test (Bonferroni-corrected for the entropy rate) is denoted by the number of asterisks (*p < 0.001, **p < 0.0001).

This was further highlighted by the significant increase in spectral entropy (Figure 8b) and entropy rate at different Markov orders (Figure 8c). The normalized LZW complexity reflects the structural richness of the signal, while the spectral entropy indicates a more uniform power distribution across frequencies, consistent with increased neural variability mainly explained by the decreased power in the alpha band (Figure 5b). The entropy rate at different orders provides a measure of the temporal predictability of the EEG signals, highlighting how psychedelics induce more diverse oscillatory patterns in the models.

## 4 Discussion

### 4.1 Main findings

Our modeling implementation of psychedelics’ effects into NMMs successfully explains the power spectrum typically observed after administration of psychedelics in the experimental literature [Carhart-Harris and Friston, 2019, Carhart-Harris and Leech, 2014, Ort et al., 2023, Timmermann et al., 2023]. Indeed, the oscillatory changes found in our personalized whole-brain models following implementation of 5-HT2AR activation closely align with the findings of Timmermann et al. (2023) [Timmermann et al., 2023], where a significant decrease in mean alpha and beta band power, together with a significant increase in mean gamma power was found in the acute phase of N,N-Dimethyltryptamine (DMT) injection in healthy subjects. Ort et al. (2023) [Ort et al., 2023] also reported similar results, showing that the mean theta and alpha power were significantly decreased in spontaneous EEG PSD under psilocybin, although no significant changes were observed in other frequencies such as the gamma band.

We found a strong correlation between the changes in mean alpha and gamma power across the brain and the density of 5-HT2ARs in each brain area. Consequently, and consistent with our initial hypothesis, the significantly reduced alpha power and increased gamma power observed under the influence of psychedelics are mainly explained in our modeling framework by the agonism of these drugs on 5-HT2A receptors and the magnitude and spatial distribution of their effects are directly related to the expression amounts of such serotonin receptors.

Our findings align with experimental evidence linking psychedelics to enhanced neural entropy and complexity [Carhart-Harris and Friston, 2019,Ruffini et al., 2023,Carhart-Harris and Leech, 2014], and support the potential therapeutic role of psychedelics in restoring the complexity deficits observed in AD [Abásolo et al., 2006, Sun et al., 2020]. Different metrics of complexity and entropy (LZW, spectral entropy, and entropy rate) resulted in increased signal diversity and richness in the personalized models under psychedelics with respect to baseline. The ability of our whole-brain models to reproduce these effects highlights their utility in exploring the neural mechanisms underlying psychedelic-induced changes and their potential for counteracting pathological states associated with reduced complexity.

### 4.2 Potential therapeutic role of psychedelics in AD

Our results suggest that psychedelics may acutely mitigate EEG spectral alterations characteristic of AD, reinforcing their therapeutic potential for this neurodegenerative condition. The spectral shifts induced by psychedelics in our models may counteract the disrupted alpha-gamma coupling and oscillatory imbalances observed in AD induced by PV dysfunction, potentially improving functional connectivity and cognition. These results indicate that psychedelics may hold therapeutic potential in the AD stages characterized by alpha power increases and reduced gamma power (prodromal and mild AD).

However, we caution that in the preclinical and very early stages of AD, cortical neurons in certain regions exhibit heightened excitability, driven partly by elevated concentrations of soluble amyloid-beta, which impairs glutamate uptake and increases extracellular glutamate levels [Zott et al., 2019, Bao et al., 2021, Harris et al., 2020]. As activation of 5-HT2ARs by psychedelics modulates glutamate’s excitatory effects, increasing the excitability of deep-layer pyramidal neurons [Andrade, 2011], their use under these conditions may intensify the existing hyperexcitability and accelerate pathological processes, suggesting a contraindication of such compounds during the earliest phases of AD. On the other hand, in the later stages of AD, the widespread loss of alpha and gamma power resulting from global neuronal death may render psychedelics ineffective as a treatment. This suggests there may be an optimal window for the treatment of AD with psychedelics, namely the prodromal to mild phase.

Importantly, the effects of psychedelics are not limited to their acute phase. Growing evidence highlights a post-acute phase characterized by enhanced neural plasticity, which underpins their long-term therapeutic effects [Ruffini et al., 2024b]. This phase involves structural and functional neural changes that can promote sustained recovery of cognitive and neural functions, which is particularly relevant for addressing the progressive degeneration seen in AD. Thus, by recalibrating neuronal excitability and oscillatory activity, psychedelics may not only acutely alleviate acute EEG abnormalities but also facilitate long-term neural reorganization and plasticity. This aligns with prior research emphasizing the role of enhanced brain functional connectivity and neuroplasticity in slowing or reversing neurodegeneration [Garcia-Romeu et al., 2022, Winkelman et al., 2023].

Finally, there is a natural synergy between psychedelics and non-invasive interventions such as transcranial electrical stimulation [Sprugnoli et al., 2021, Dhaynaut et al., 2022] or sensory stimulation [Iaccarino et al., 2016, Martorell et al., 2019, Adaikkan et al., 2019, Adaikkan and Tsai, 2020], which seek to entrain natural fast frequency circuits in AD. The combination of psychedelics with such interventions can produce benefits from the acute effects of each and by the increased window of plasticity facilitated by psychedelics [Ruffini et al., 2024b, Ruffini et al., 2024a].

### 4.3 Limitations and future work

Several limitations in the present study are worth pointing out. Although our focus is on AD, the inclusion of data from healthy subjects could offer a critical comparative framework. By contrasting the altered alpha and gamma power in AD models treated with psychedelics against the normative power spectrum dynamics observed in healthy individuals, we might better understand the restorative potential of psychedelics. Such comparisons could elucidate whether the modeled oscillatory dynamics approximate—or fully restore—the canonical signatures of healthy brain activity.

Furthermore, our AD model represents pathology only at the level of global connectivity and external noise. In parallel work in mesoscale modeling of AD, we have developed an adapted version of the LaNMM that reproduces the impact of PV synapse damage in oscillatory dynamics, producing slowing of EEG, increased alpha power, and reduced gamma [Sanchez-Todo et al., 2024]. In future work, we can investigate the effects of psychedelics in this AD mesoscale model, which incorporates physiologically-inspired mechanisms to reproduce the disruptions in oscillatory activity observed in AD.

## 5 Conclusions

In this study, we explored the mechanisms behind spectral power alterations during the acute phase of serotonergic psychedelic drug effects and provided a novel mechanistic understanding of their impact on brain oscillatory dynamics. By implementing 5-HT2A receptor activation in a laminar neural mass model, we reproduced the characteristic EEG multiband power spectrum alterations observed under psychedelics within a single cortical column. Extending this approach, we used 30 subject-specific whole-brain models, integrating MRI, dMRI, and fMRI data from AD subjects, to simulate the effects of psychedelics on whole-brain dynamics following 5-HT2A receptor stimulation.

The subject-specific whole-brain models effectively reproduced the power spectrum changes observed experimentally with psychedelics, notably the significant decreases in alpha and beta power and increases in gamma power. Our findings align with previous studies combining fMRI and EEG approaches to psychedelics [Timmermann et al., 2023, Ort et al., 2023] and further demonstrate a strong correlation between modulation of mean alpha and gamma power across brain regions and the spatial distribution of 5-HT2A receptors. By combining these whole-brain models with a biophysical head model, we successfully reproduced the suppression of alpha rhythms and increased gamma power observed in prior EEG studies as well as the complexity changes associated with the effects of psychedelics.

This modeling framework offers new insights into the neural mechanisms underlying psychedelic action and highlights the potential of psychedelics to restore healthy oscillatory dynamics in disorders like AD. Our approach paves the way for using computational whole-brain models to investigate the potential of psychedelics in rebalancing neural activity in neurodegenerative and psychiatric disorders.

## Data availability

Data used in the preparation of this article were obtained from the Alzheimer’s Disease Neuroimaging Initiative (ADNI) database (http://adni.loni.usc.edu). The Alzheimer’s Disease Neuroimaging Initiative (ADNI) is a longitudinal, multicenter study aimed at developing clinical, imaging, genetic, and biochemical biomarkers for the early detection and tracking of Alzheimer’s disease (AD). A key feature of ADNI is its open data-sharing policy, which allows qualified researchers worldwide to access a comprehensive collection of de-identified data, including structural, functional, and molecular brain imaging, biofluid biomarkers, cognitive assessments, genetic data, and demographic information.

## Acknowledgements

We want to thank the Elite Master Program in Neuroengineering at the Technical University of Munich, funded by the Elite Network of Bavaria, for the financial support that facilitated the publication of this work.

## Contributions

Conceptualization: JCG, EL-S, FC, RS-T, JV, RGA, GR. Formal analysis: JCG, EL-S. Funding acquisition: JCG, GR. Investigation: JCG. Methodology: JCG, EL-S, EL-C, RS, GR. Software: JCG, EL-S, EL-C, RS, GR. Visualization: JCG, EL-S, RS-T. Writing – original draft: JCG, EL-S. Writing – review & editing: JCG, EL-S, FC, EL-C, RS-T, RS, JV, RGA, GR.

## Supplementary Information

### S1 Potential therapeutic effects of psychedelics in AD Stages of AD and electrophysiological alterations

AD progresses through distinct stages, beginning with a *preclinical* phase, characterized by pathological changes such as amyloid-beta and tau accumulation detectable through biomarkers, but without noticeable cognitive symptoms [Sperling et al., 2011]. The next stage, *prodromal AD* or Mild Cognitive Impairment (MCI) due to AD, involves subtle cognitive declines, particularly in memory, that exceed normal aging but do not yet significantly impair daily life [Petersen, 2004]. The disease progresses then from *mild AD dementia*, where cognitive impairments interfere with daily tasks, to *moderate AD dementia* with severe cognitive decline and increased dependency, and ultimately to *severe AD dementia* marked by extensive neurodegeneration [Alzheimer’s, 2020].

In the preclinical stages of AD, some studies report an increase in alpha power along with a potential increase in gamma power [Gaubert et al., 2019,Nakamura et al., 2018]. However, findings regarding gamma power in this stage are variable, with some evidence suggesting stable or mildly reduced gamma activity [Rochart et al., 2020].

After the preclinical stage of AD, the EEG spectrum undergoes a characteristic slowing, with increased power in lower frequencies (delta and theta) and decreased power in higher frequencies (beta and gamma) as the disease progresses [Casula et al., 2022,Babiloni et al., 2020, Murty et al., 2021]. Increased alpha power and hypersynchronization in alpha-band functional connectivity reflect an imbalance in excitation and inhibition, which contributes to neuronal hyperactivity and network disruptions [Nakamura et al., 2018, López et al., 2014], although there is also evidence of decreased alpha power in MCI patients [López-Sanz et al., 2016].

As the disease advances into early to moderate AD, alpha power begins to decline, and a slowing of the alpha peak frequency is observed, which correlates with structural and functional impairments [Passero et al., 1995, Garcés et al., 2013, Puttaert et al., 2021]. Gamma oscillations, which are essential for higher-order cognitive functions like working memory and sensory processing, show a more consistent decrease during MCI and early AD, reflecting synaptic dysfunction and impaired connectivity [Babiloni et al., 2020, Sanchez-Todo et al., 2024, Murty et al., 2021].

In later stages of AD, both alpha and gamma power are markedly reduced, alongside significant increases in slower rhythms such as delta and theta. This shift in the EEG power spectrum mirrors widespread cortical atrophy, disrupted excitatory-inhibitory balance, and breakdowns in network synchronization and efficiency, which underpin the severity of cognitive and functional decline [Casula et al., 2022, Babiloni et al., 2020].

#### Physiopathology of AD

AD is associated with the dysfunction of inhibitory interneurons, particularly those expressing parvalbumin (PV), and the resulting network abnormalities in AD [Palop and Mucke, 2016, Sanchez-Todo et al., 2024]. PV interneurons, crucial for maintaining fast oscillations, become impaired in AD due to both direct pathology and, possibly, compensatory neural changes [Palop and Mucke, 2016]. This dysfunction leads to a loss of gamma rhythm, essential for higher cognitive processes, and contributes to network instability and hypersynchrony in slower frequencies.

In early AD, synaptic connections from PV interneurons in superficial layers—critical for generating fast gamma rhythms—are among the first to be disrupted. This early damage corresponds with the emergence of increased alpha and reduced gamma activity observed in MCI and mild-to-moderate AD. Eventually, persistent alpha hypersynchronization further deteriorates these circuits and affects other connected areas in later disease stages. While the precise mechanisms linking PV interneuron dysfunction to EEG slowing remain unclear, and compensatory processes have been suggested [Gaubert et al., 2019], a modified version of LaNMM demonstrates how disrupting PV synapse connectivity can reproduce these electrophysiological changes [Sanchez-Todo et al., 2024]. In this model, slowing of the EEG emerges from the interaction between disrupted fast circuits and slow circuits rather than compensation.

#### Restoring healthy oscillatory balance

Recent research suggests that restoring PV interneuron function or modulating network synchrony could help re-establish healthy oscillatory dynamics and improve cognitive function in AD, which motivates the use of gamma stimulation in AD, for instance, through sensory stimuli or transcranial alternating current stimulation (tACS) [Palop and Mucke, 2016, Iaccarino et al., 2016, Martorell et al., 2019, Fatemi et al., 2021, Sanchez-Todo et al., 2024]. Building on this idea, psychedelics could restore the oscillatory balance in AD, especially in the early stages of the disease. Psychedelics reduce alpha power, enhancing network flexibility and interconnectivity [Nutt et al., 2020, Carhart-Harris and Friston, 2019], and may increase gamma power, promoting global connectivity and enhancing oscillations associated with cognitive processing [Silverstein et al., 2024, Kometer et al., 2013, Muthukumaraswamy et al., 2013, Smausz et al., 2022]. By modulating 5-HT2A receptors on cortical pyramidal cells, these compounds may indirectly engage PV interneurons, promoting gamma synchrony and balancing excitatory-inhibitory networks [Halberstadt, 2015]. Gamma power increases through psychedelics could potentially counteract the reduction observed in AD, further aiding in the restoration of cognitive functions that depend on these rhythms. Psychedelics also reduce delta and theta activity, counteracting the increased power in these bands often observed in AD [Pallavicini et al., 2019]. This modulation of slow oscillations may help attenuate pathological hypersynchrony in delta and theta bands, restoring a healthier balance across frequencies that aligns with normal cognitive processing.

The enhanced excitation under psychedelics is also related to increased neural entropy and complexity of ongoing brain activity, as reported extensively in the literature [Carhart-Harris and Friston, 2019, Ruffini et al., 2023], which could counteract the reduction in complexity observed in AD subjects [Abásolo et al., 2006, Sun et al., 2020]. Psychedelics also modulate network connectivity patterns, increasing global integration and decreasing modularity [Tagliazucchi et al., 2016]. This shift could address the network decoupling seen in AD, promoting a healthier, more flexible connectivity pattern. The combined effects on gamma power, slow oscillations, and network integration suggest that psychedelics may help re-establish synchronous activity and alleviate cognitive deficits through PV interneuron engagement, oscillatory enhancement, and network stabilization. This rationale motivates the study of psychedelics as potential modulators of oscillatory dynamics and connectivity, addressing key electrophysiological deficits associated with disease progression and cognitive impairment.

### S2 LaNMM description

#### S2.1 Single LaNMM equations and parameters

The LaNMM [Sanchez-Todo and Bastos, 2023] employs a second-order differential equation to model the average membrane potential perturbation in a neuronal population *m* caused by inputs from another population *n*. This equation captures the synaptic dynamics within the average neuron of a given population, translating the presynaptic mean firing rate φ_*n*_ into a perturbation of the mean membrane potential *u*_*m*←*n*_ in the postsynaptic population *m*. This relationship is expressed through the integral operator 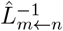, defined as:

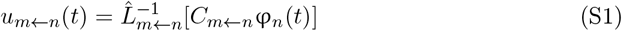

where *C*_*m*←*n*_ is the connectivity constant, indicating the average number of synaptic contacts between population types, which inverse is the differential operator 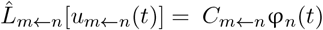, defined as

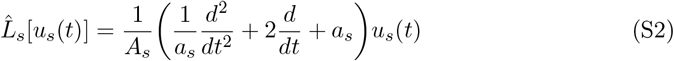

where *A*_*s*_ is the average excitatory/inhibitory synaptic gain and *a*_*s*_ is the rate constant of the connection (*a*_*s*_ = 1*/τ*_*s*_, *τ*_*s*_ denoting the synaptic time constant). Note that, for simplicity, the single index *s* is used to represent each average synapse (i.e., connection) from one neuronal population to another, so that *s* ≡ {*m* ← *n* : *C*_*m*←*n*_ ≠ 0 } . In the LaNMM, *s* ∈ [1, 13] and the neural populations are denoted by (*m, n*) ∈ [*P*_1_, SS, SST, *P*_2_, PV, *e*1, *e*2]. The indices assigned to each connection in the LaNMM are depicted in Fig. S1. Finally, the average output firing rate φ_*m*_ of population *m* is defined by the sigmoid function

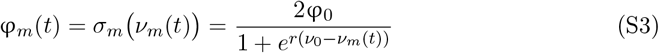

where φ_0_ is half of the maximum firing rate of the population, *ν*_0_ the value of the potential when the firing rate is φ_0_ and *r* determines the slope of the sigmoid at the central symmetry point (*ν*_0_,φ_0_). Here, *ν*_*m*_ denotes the average membrane potential of the postsynaptic population *m*

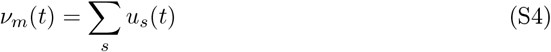

where *u*_*s*_ is the membrane perturbation per each of its incoming connections *s*. Consequently, the general differential equation capturing the average synaptic dynamics of each connection *s* in the LaNMM between each postsynaptic and presynaptic population pair *m* ← *n* is defined as

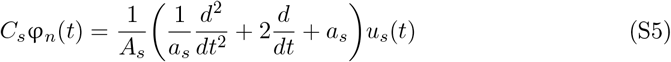

which depends on the average membrane potential perturbation caused by each connection *u*_*s*_ inputting the presynaptic population *n*, and the average membrane potential *ν*_*n*_ and average firing rate φ_*n*_ of this presynaptic population.

Notice that this modeling approach focuses on the dynamics of the connections between neural populations instead of the populations’ dynamics, as opposed to the traditional approach followed in neural mass modeling. This is referred to by Sanchez-Todo et al. (2023) [Sanchez-Todo and Bastos, 2023] as the *synapse-driven* formulation of an NMM, which simplifies the definition of the neural dynamics generalizing the model equations. The equation for each connection describing the LaNMM is shown hereafter:

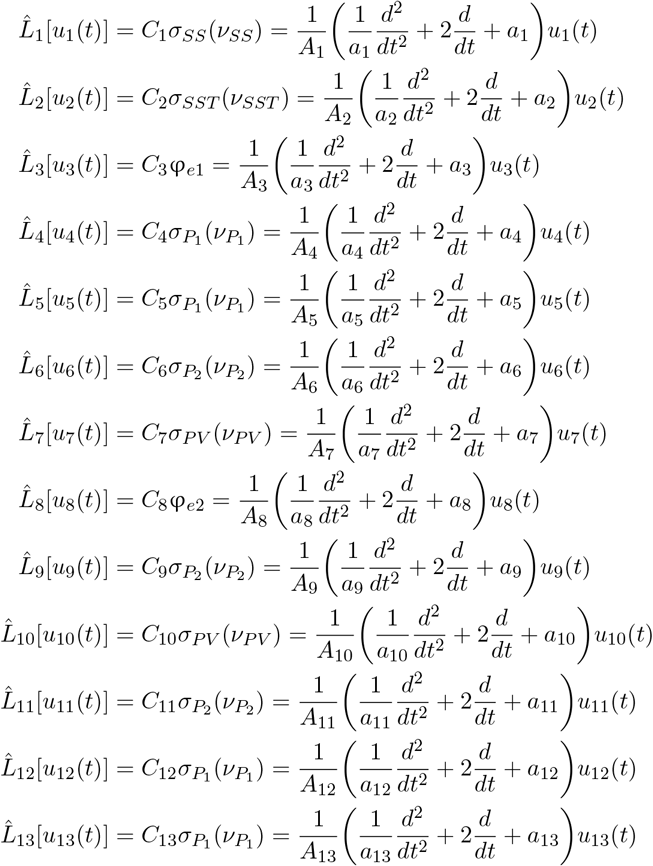

with the average membrane potential *ν*_*m*_ of each neural population in the LaNMM given by

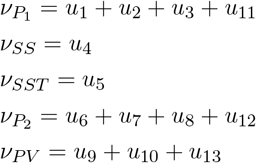

The list of LaNMM parameters used for the simulations is shown in Table S1. Model parameters were adopted from Sanchez-Todo et al. [Sanchez-Todo and Bastos, 2023], except for the standard deviation *σ*_*e*_ of the external noisy perturbations *e*_1_ and *e*_2_. In the single LaNMM study, we adjusted *σ*_*e*_ to match the average SD value of the external inputs received by each LaNMM in the whole-brain model personalization (see Section S2.3).

**Figure S1:**
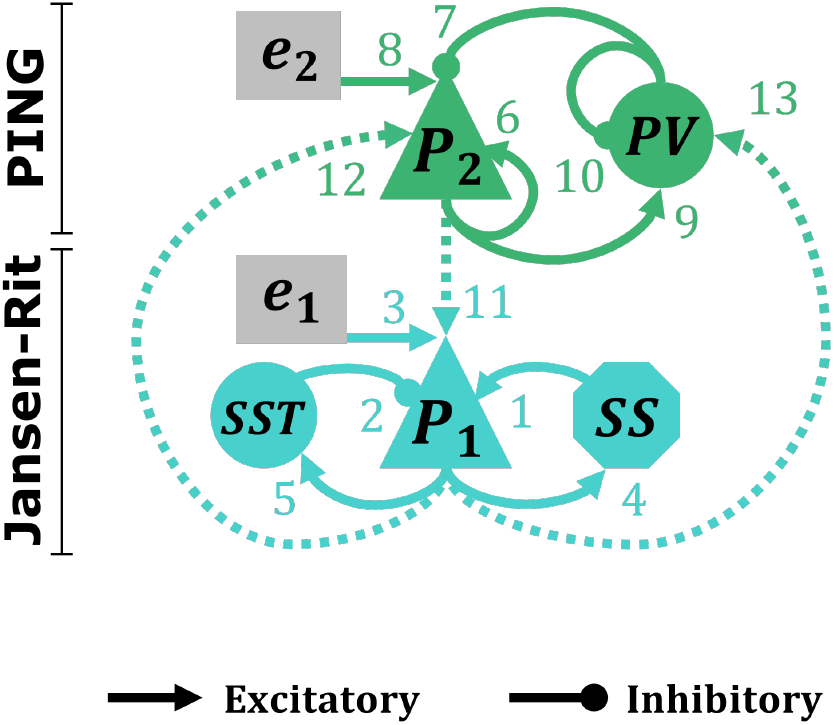
Schematic diagram of the LaNMM depicting its neuronal populations and the connections between them. The PING sub-circuit is depicted in green and the JR sub- circuit in blue. Rounded shapes indicate inhibitory populations and interactions, triangles and octagons indicate excitatory ones, and squares represent external inputs. Numbers indicate the index *s* assigned to each connection. Dashed lines represent connections between the PING and Jansen-Rit parts sub-circuits. The neural population abbreviations (*P*_1_, SS, SST, *P*_2_, PV, *e*1, *e*2) are defined in the main text.

#### S2.2 Long-range LaNMM connectivity

The long-range connections between nodes in the model are weighted following the connectivity patterns between cortical regions described by Bastos et al. [Bastos et al., 2012] and recent modeling work [Mejias et al., 2016, Jaramillo et al., 2019]. In particular, *P*_1_ and *P*_2_ populations of different parcels *i, j* are connected as follows: 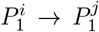 (lateral connection), 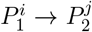 (feedback or descending connection) and 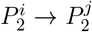 (feedforward or ascending connection) with relative weights *w* of 0.5, 0.5 and 1 respectively (see Figure S2). The resulting connectivity *C*_*ij*_ between parcels *i* and *j* depends on which of these three connections is involved, and is computed as:

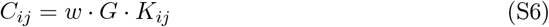

*w* represents the relative strength of the connection, *G* is the global scaling factor that adjusts the overall connection strength and *K*_*ij*_ is the connectome value. For simplicity, we assumed that all long-range cortico-cortical connections across brain columns are mediated via pyramidal populations — hence, they all have an excitatory effect.

#### S2.3 Whole-brain model external inputs

The external noisy inputs *e*1 and *e*2 applied to the JR and PING model’s pyramidal populations (see Section 2.1) are modeled in the whole-brain models as the sum of two additive noises:

- On the one hand, a *common noise* is applied homogeneously to all parcels (i.e., the same realization of noise is applied to every parcel) and modeled as white Gaussian noise with mean *µ*_*c*1_ = 270 Hz to *P*_1_ and mean *µ*_*c*2_ = 90 Hz to *P*_2_, mirroring the values of the single LaNMM [Sanchez-Todo and Bastos, 2023]. The SD of the common noise *σ*_*c*_ is personalized for every subject.
- On top of this, white Gaussian noise with mean *µ*_*i*_ = 0 Hz and SD *σ*_*i*_ = 20 Hz is added to the common noise in both *P*_1_ and *P*_2_ *homotopically*, i.e., the same realization of noise is applied to “mirror” areas of homologous parcels in both hemispheres—e.g., same noise realization to the left and right insula.

**Table S1:** Parameters, description, and values of the baseline Laminar NMM employed in this work. The values are taken from Sanchez-Todo et al. [Sanchez-Todo and Bastos, 2023], except for the external inputs (φ_*e*1_, φ_*e*2_). Notice that the sigmoid function parameters defining the firing rate φ_*m*_ in this model (*ν*_0_, φ_0_, *r*), apply to all neuron populations. Moreover, all excitatory connections follow the same synapse dynamics, (*A, a*)_AMPA_ = (*A, a*)_1,3,4,5,6,8,9,11,12,13_ while all inhibitory connections follow either fast, 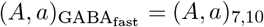, or slow dynamics, 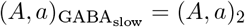.

| Parameter | Description | Value |
| --- | --- | --- |
| $A_s$ | Average excitatory and (fast and slow) inhibitory synaptic gains | $A_{\text{AMPA}} = 3.25$ mV,<br>$A_{\text{GABA}_{\text{fast}}} = -22$ mV,<br>$A_{\text{GABA}_{\text{slow}}} = -30$ mV |
| $a_s$ | Time constant of average excitatory and inhibitory postsynaptic potentials | $a_{\text{AMPA}} = 100$ Hz,<br>$a_{\text{GABA}_{\text{fast}}} = 50$ Hz,<br>$a_{\text{GABA}_{\text{slow}}} = 220$ Hz |
| $C_s$ | Connectivity constant between populations | $C_1 = 108, C_2 = 33.75,$<br>$C_3 = 1, C_4 = 135,$<br>$C_5 = 33.75, C_6 = 70,$<br>$C_7 = 550, C_8 = 1,$<br>$C_9 = 200, C_{10} = 100,$<br>$C_{11} = 80, C_{12} = 200,$<br>$C_{13} = 30$ |
| $\nu_0$ | Potential at 50% of the firing rate | 6 mV<br>(except $P_2$'s $\nu_0 = 1$ mV) |
| $\varphi_0$ | Half of the maximum firing rate | 2.5 Hz |
| $r$ | Slope of the sigmoid function at $\nu_0$ | $0.56 \text{ mV}^{-1}$ |
| $\mu_e, \sigma_e$ | Mean and SD of the external input firing rates $\varphi_e$ | $\varphi_{e1}$: white noise with<br>$\mu_{e1} = 270$ Hz and $\sigma_{e1} = 34$ Hz<br>$\varphi_{e2}$: white noise with<br>$\mu_{e2} = 90$ Hz and $\sigma_{e2} = 34$ Hz |

**Figure S2:**
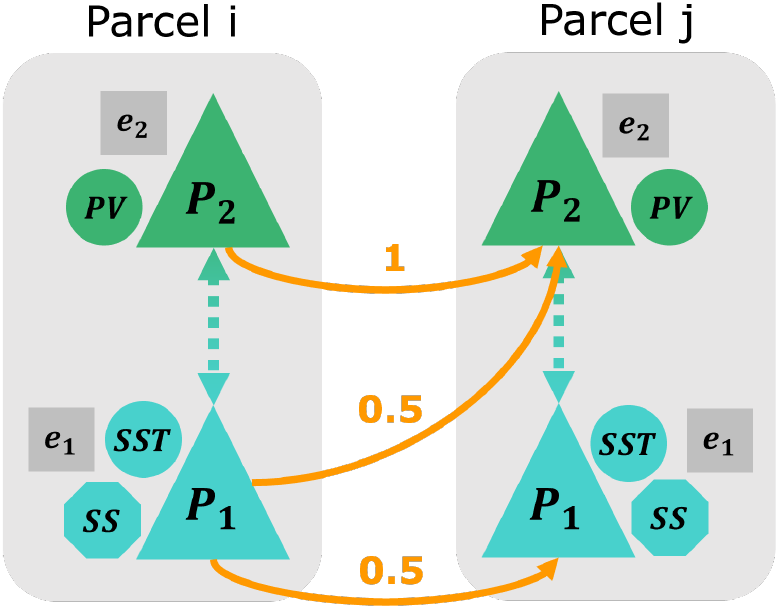
Long-range cortico-cortical parcel connections and their relative strength. Note that, for simplicity, cortico-cortical connections are only drawn in one direction, but they are present in both directions in the model.

These two external inputs model the influence of FC pathways involving subcortical structures such as the thalamus [Uddin et al., 2008, Wang et al., 2019], constituting a rough thalamo-cortical model, which may play a pivotal role in the context of psychedelics [Müller et al., 2017].

### S3 Neuroimaging data acquisition parameters

The model personalization pipeline involves three types of neuroimaging data: MRI, dMRI, and blood-oxigen-level-dependent (BOLD) resting-state fMRI (rs-fMRI). The data acquisition parameters for these modalities and the patient cohort selected is provided below.

T1-weighted MRI images were acquired using standardized MPRAGE protocols with 3T MRI scanners across multiple sites. Key acquisition parameters included TR = 2300 ms, TE = 2.98 ms, voxel size = 1×1×1 mm, and flip angle = 9°. Resting-state fMRI data were acquired using standardized ADNI3 protocols with gradient-echo EPI sequences. Key parameters included TR = 3000 ms, TE = 30 ms, flip angle = 90°, voxel size = 3.3×3.3×3.3 mm, and scan duration = ∼6 minutes. Diffusion MRI was acquired with b-values of 0 and 1000 s/mm^2^, 41 gradient directions, TR = ∼9100 ms, TE = ∼98 ms, and voxel size = 2.7×2.7×2.7 mm. Detailed imaging protocols and scanner-specific adjustments are available in the ADNI3 documentation.

### S4 Parcellation of PET-derived 5-HT2AR density in DK-68

To obtain a quantitative measure of the 5-HT2AR density in each DK-68 parcel, we parcellated Beliveau et al.’s 5-HT2AR average density map into the DK-68 cortical atlas. For this purpose, we used the *Neuromaps* toolbox [Markello et al., 2022] and followed Hansen et al.’s parcellation pipeline [Hansen et al., 2022]. We used the DK-68 average subject template atlas automatically labeled by FreeSurfer 7.4 in *fsaverage* space [Fischl, 2012, Fischl et al., 2004] to obtain the average 5-HT2AR density in the surface area of each cortical parcel. The PET imaging data used in this study, including receptor images and densities, are included in Neuromaps^1^ and the DK-68 *fsaverage* atlas is available in the installation files of *FreeSurfer* 7.4^2^. The obtained receptor map and the DK-68 parcellation used can be found in Figure S3.

**Figure S3:**
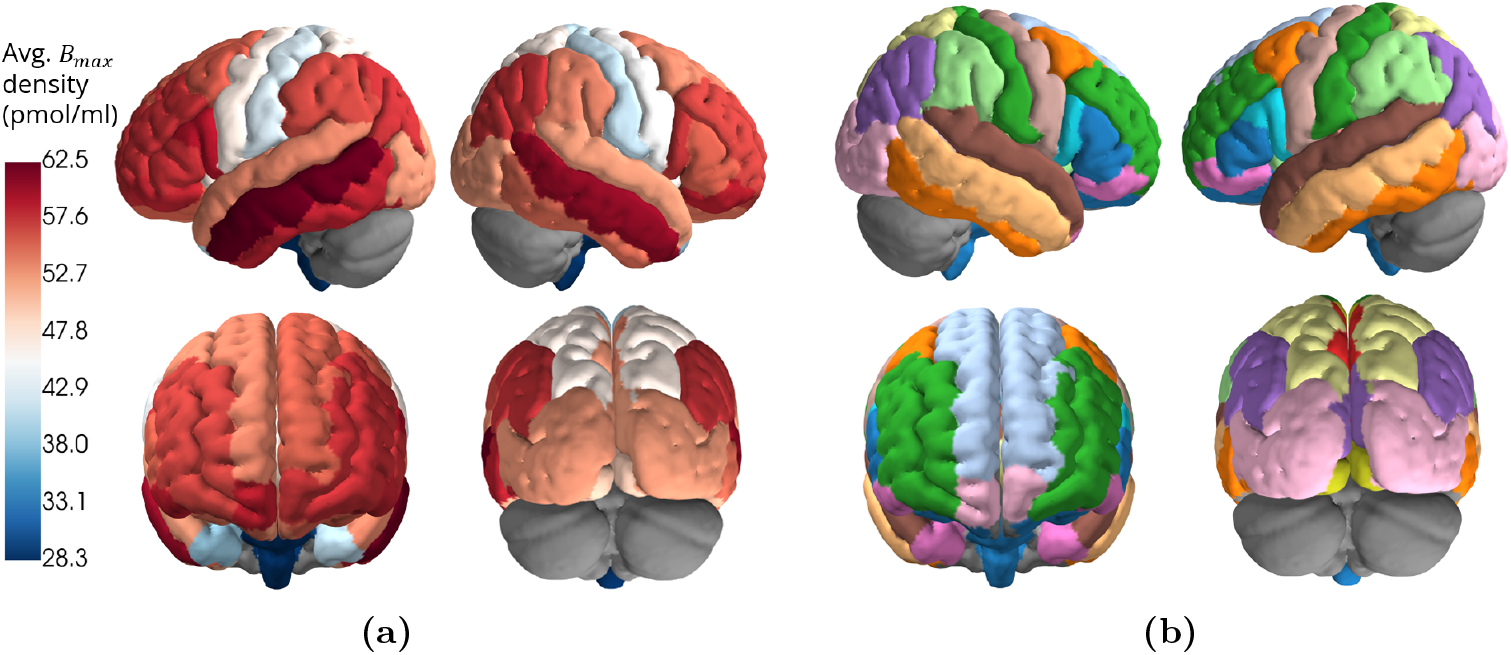
**(a)** 5-HT2A receptor map of average *B*_*max*_ densities parcellated into the DK- 68 atlas. **(b)** DK-68 atlas displaying each parcel in distinct colors, with homotopic parcels sharing the same color. Excluded areas (i.e., subcortical structures) are displayed in grey.

### S5 Biophysical head model generation

Figure S4 illustrates the process of creating the template head model. The head model was generated from a template T1-weighted image (MNI-ICBM 152 non-linear 2009^3^). Segmentation was performed with a pre-trained convolutional neural network (based on a stacked 2D U-Net [Ronneberger et al., 2015]), generating volume masks for the scalp, skull, CSF (including the ventricles), GM and WM. All segmentation masks were manually inspected and corrected as needed. The volume segmentations were then converted to triangulated surface meshes and then to a tetrahedral volume mesh using the SimNIBS library v3.2.3^4^.

The lead field matrix was then calculated for all electrode positions defined in the 10- 10 EEG positioning system [Jurcak et al., 2007]. This is a *N*_mesh nodes_ × *N*_electrodes_ − 1 matrix where each column contains the *E*-field component orthogonal (*E*_*n*_) to the cortical surface (GM-CSF interface) induced by a bipolar combination of electrodes where Cz is always defined as the cathode (-1 mA). Electrodes were modeled as a cylinder of gel with a height of 3 mm and a radius of 1 cm (representing Neuroelectrics’ NG-Pistims^5^). E-field calculations were performed with the SimNIBS library, assigning to each head tissue an isotropic and homogeneous conductivity: 0.33 S/m, 0.008 S/m, 1.79 S/m, 0.4 S/m and 0.15 S/m, respectively for scalp, skull, CSF (including the ventricles), GM and WM. The gel representing the electrodes was modeled with a conductivity of 4.0 S/m.

The lead-field matrix was then used to map activity from source space to electrodes using the reciprocity theorem as described in Ruffini (2015) [Ruffini, 2015]. This was performed by first mapping source activity per parcel to activity per mesh node. To do so, initially the activity of a parcel was assigned to all the mesh nodes in the parcel. This initial map was then blurred by applying a linear operator (**B**) that changes the activity per node to a weighted average of the activity of neighboring nodes, where the weights are determined based on the geodesic distance between nodes and the area of each node of the mesh (defined as the sum of the areas of all the triangles of the mesh that share the node, divided by 3). The electrode voltage differences were then calculated by performing the matrix multiplication of the lead-field matrix and the cortical map of the sources.

**Figure S4:**
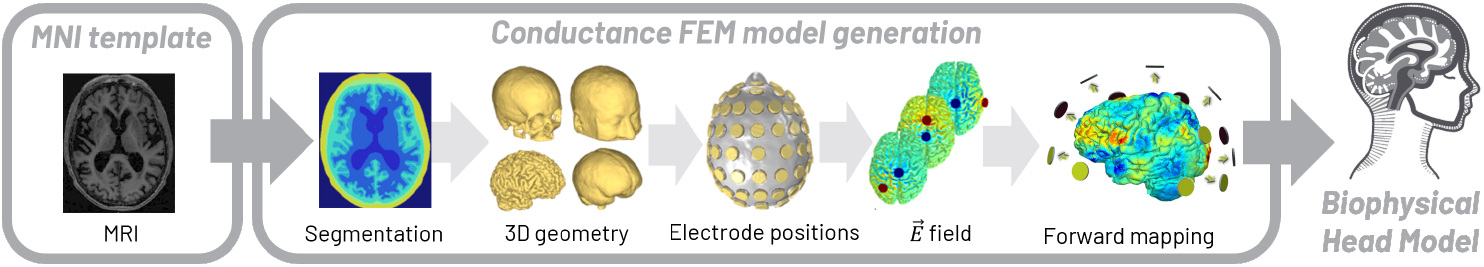
Head model template and cortical mapper generation pipeline. Figure adapted from Salvador et al. (2022) [Salvador et al., 2022]

### S6 Uncoupled LaNMM oscillatory behavior

We studied the frequency with the highest PSD (i.e., dominant frequency) in the pyramidal populations over *A*_*L*5*P*_ ∈ [0, 10] mV, as well as the changes in the membrane potential of *P*_1_ and *P*_2_ resulting from changing the *A*_*L*5*P*_ parameter (Figure S5).

**Figure S5:**
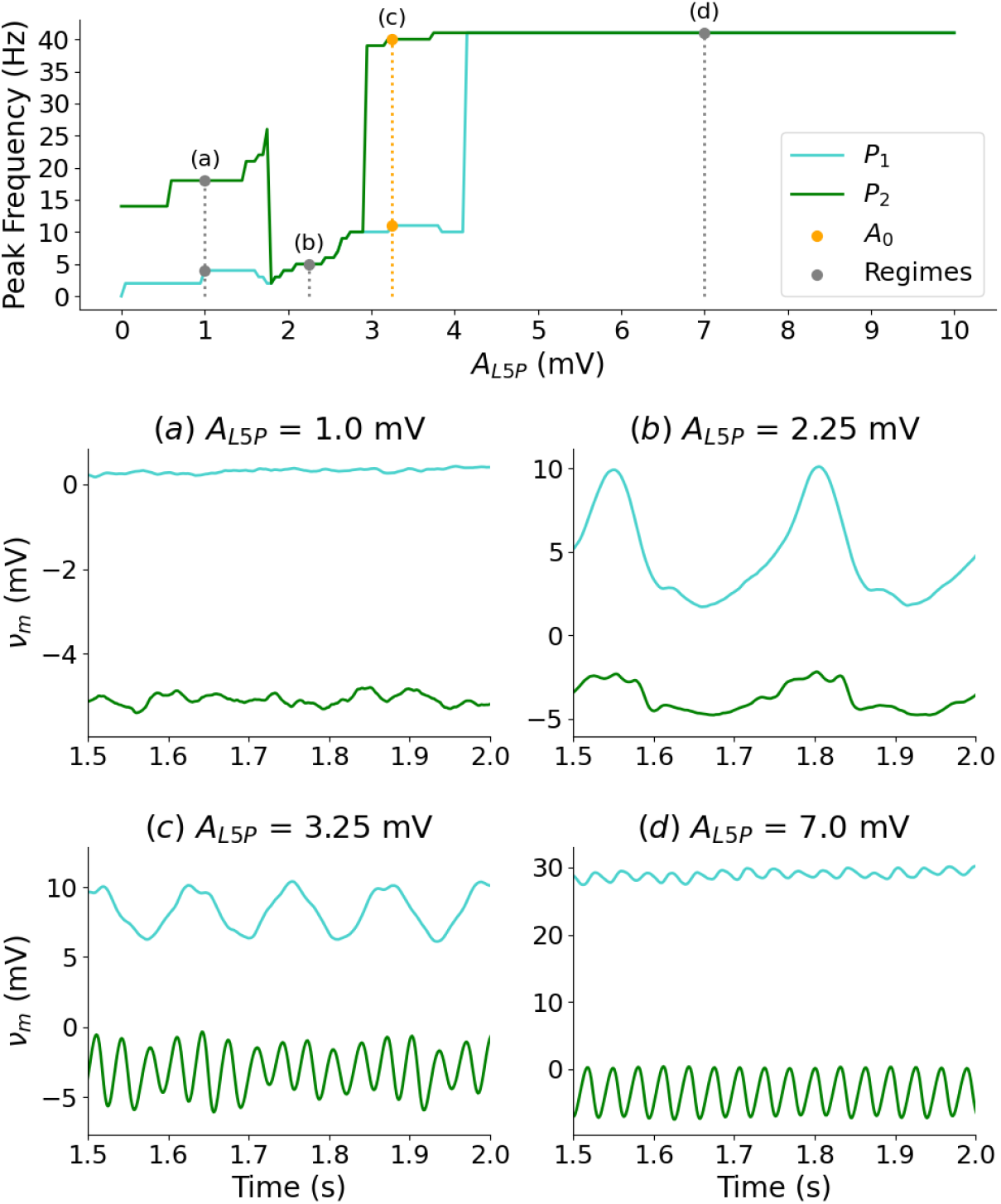
**Top:** Dominant frequency of the membrane potentials of *P*_1_ (blue) and *P*_2_ (green) pyramidal populations in an uncoupled LaNMM for different values of the synaptic gain *A*_*L*5*P*_ (averaged across five realizations). The orange dashed line denotes the pyramidal gain at baseline *A*_0_ and the grey points denote different model regimes. **Bottom:** Membrane potential of *P*_1_ (blue) and *P*_2_ (green) populations for different *A*_*L*5*P*_ values. The plotted *A*_*L*5*P*_ value is shown on top of each figure. (a) No oscillations regime. (b) Theta/delta oscillations regime. (c) Alpha and gamma oscillations regime. (d) Gamma oscillations regime.

We identify four different dynamical regimes in Figure S5:

1. For *A*_*L*5*P*_ ∈ [0, 1.8) mV (Figure S5a), no oscillations are present in any pyramidal population (noisy regime).
2. For *A*_*L*5*P*_ ∈ [1.8, 2.75) mV (Figure S5b), theta and delta oscillations are dominant in both pyramidal populations.
3. For *A*_*L*5*P*_ ∈ [2.75, 4.15) mV (Figure S5c), alpha oscillations are dominant in *P*_1_ and gamma oscillations in *P*_2_ (LaNMM’s default regime).
4. For *A*_*L*5*P*_ ∈ [4.15, ∞) mV (Figure S5d), gamma oscillations are dominant in both pyramidal populations.

To assess whether the spectral power alterations in our model are frequency-specific, we studied the power spectrum of the pyramidal populations over the synaptic gain *A*_*L*5*P*_ ∈ [0, 10] mV and applied a square window for frequencies *f* = [0, 100] Hz (Figure S6). The findings show that the area with the highest PSD in *P*_1_ aligns with the delta-theta band for *A*_*L*5*P*_ ∈ [2, 2.75] mV, the alpha band for *A*_*L*5*P*_ ∈ [2.75, 3.75] mV, and gamma band for *A*_*L*5*P*_ *>* 3.75 mV. In population *P*_2_, this area lies within the gamma band and agrees with gain values related to gamma oscillations in the previous section. There is also increased PSD in the beta band for both populations, associated with the lower-power harmonics of the alpha band. We conclude that the most significant changes induced by psychedelics in the PSD magnitude in our LaNMM are specific to the alpha and gamma bands within the gain range under study [*A*_0_, *A*_max_] in both *P*_1_ and *P*_2_ populations.

**Figure S6:**
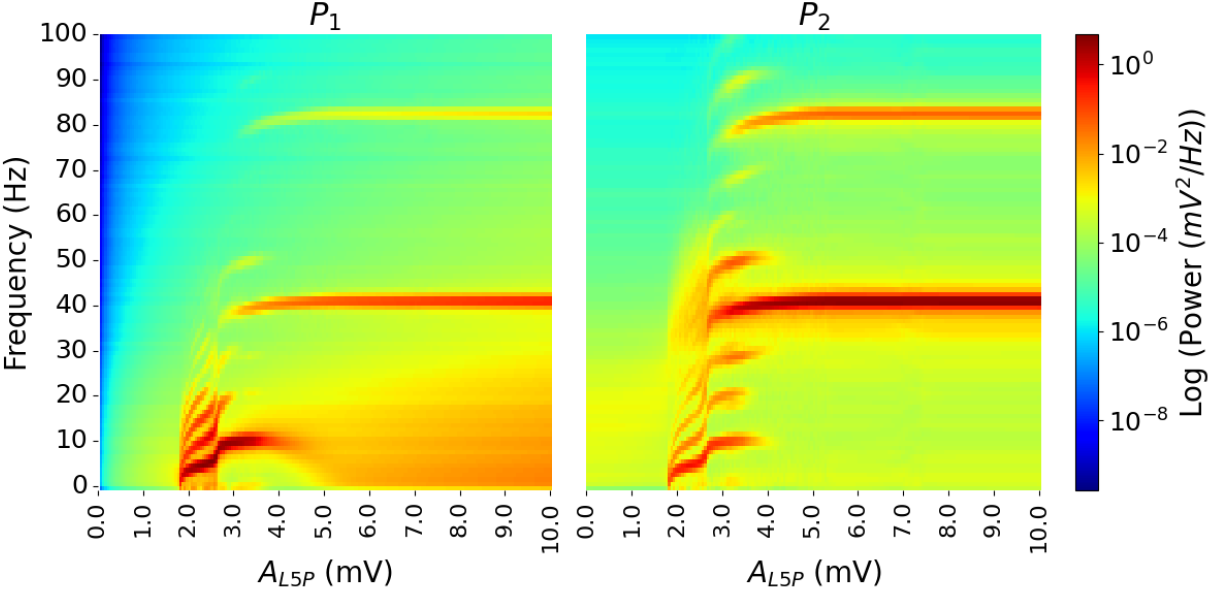
Power spectrum of *P*_1_ (left) and *P*_2_ (right) populations membrane potential over *A*_*L*5*P*_ parameter. The PSD (color scale) is shown in a logarithmic scale.

### S7 Whole-brain model fitting results

Figures S7 and S8 show the values of the optimized parameters *G* and *σ*_*c*_*/σ*_*i*_ for each subject after model personalization.

**Figure S7:**
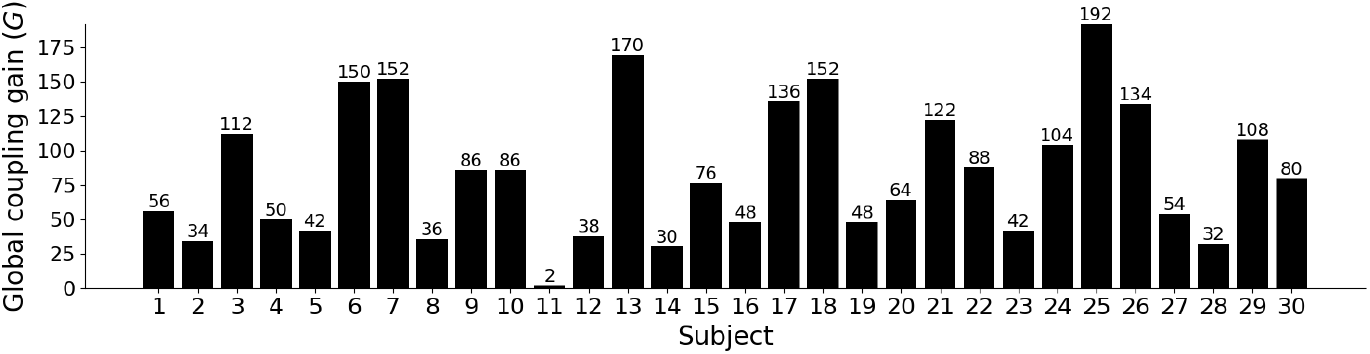
Personalized values of the *G* parameter after model personalization

**Figure S8:**
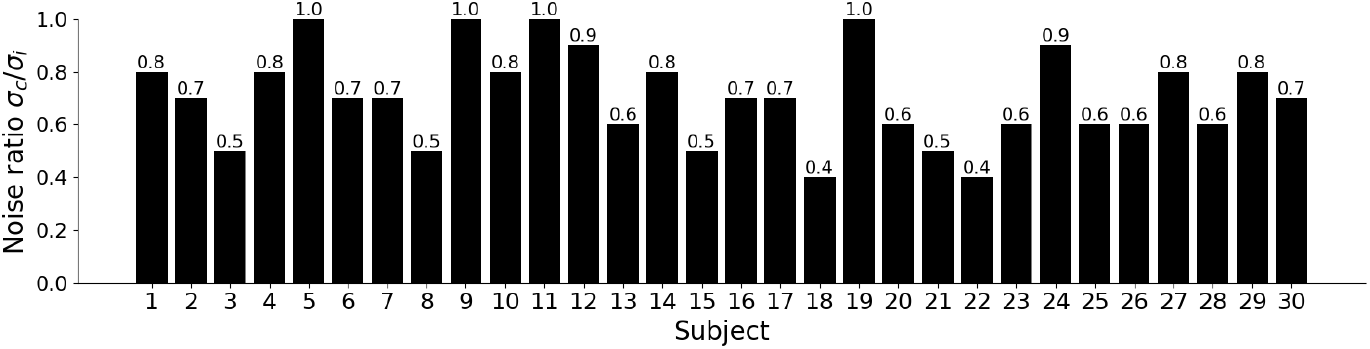
Personalized values of *σ*_*c*_*/σ*_*i*_ after model personalization

Figure S9 shows the FC matrices obtained from the synthetic BOLD signal for the optimal parameter selection for each subject, compared with the subject’s empirical BOLD FC. The optimal value of G and the resulting value of PCC and RMSE between empirical and synthetic matrices is also displayed.

**Figure S9:**
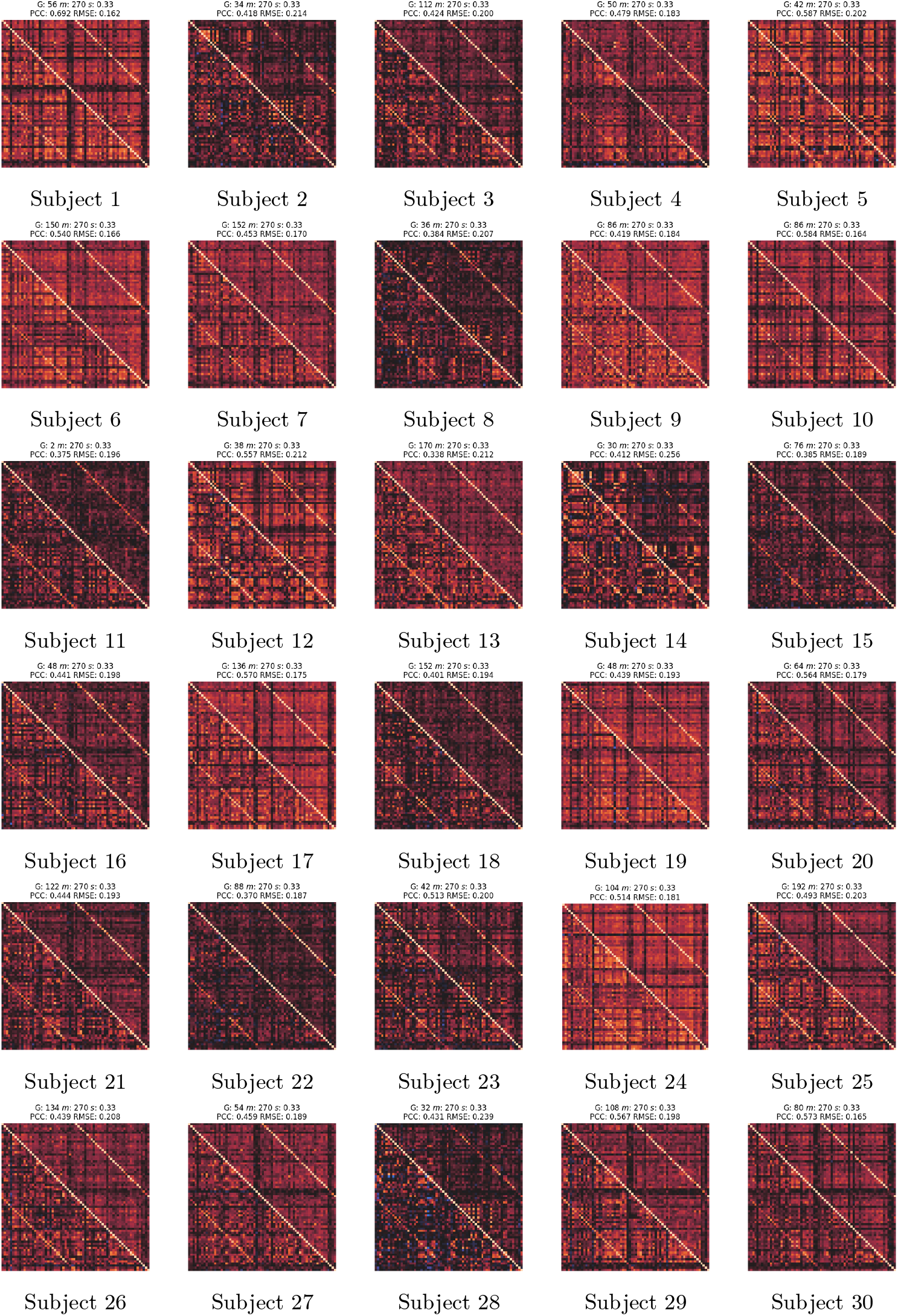
Comparison between the empirical rs-fMRI BOLD FC (lower triangular matrix) and the synthetic FC obtained with the fitted model (upper triangular matrix) for the best set of parameters selected for each subject.

## Footnotes

1 https://github.com/netneurolab/neuromaps

2 http://surfer.nmr.mgh.harvard.edu

3 https://nist.mni.mcgill.ca/atlases/

4 https://github.com/simnibs/simnibs

5 https://www.neuroelectrics.com/solution/spareparts-consumables/ngpistim

## References

Abásolo, D., Hornero, R., Espino, P., Alvarez, D., and Poza, J. (2006). Entropy analysis of the EEG background activity in Alzheimer’s disease patients. Physiological Measurement, 27(3):241–253.

Adaikkan, C., Middleton, S. J., Marco, A., Pao, P.-C., Mathys, H., Kim, D. N.-W., Gao, F., Young, J. Z., Suk, H.-J., Boyden, E. S., McHugh, T. J., and Tsai, L.-H. (2019). Gamma Entrainment Binds Higher-Order Brain Regions and Offers Neuroprotection. Neuron, 102(5):929–943.e8.

Adaikkan, C. and Tsai, L.-H. (2020). Gamma Entrainment: Impact on Neurocircuits, Glia, and Therapeutic Opportunities. Trends in Neurosciences, 43(1):24–41.

Alzheimer’s, A. (2020). 2020 Alzheimer’s disease facts and figures. Alzheimer’s & Dementia: The Journal of the Alzheimer’s Association.

Andrade, R. (2011). Serotonergic regulation of neuronal excitability in the prefrontal cortex. Neuropharmacology, 61(3):382–386.

Avants, B., Tustison, N., and Song, G. (2009). Advanced normalization tools (ANTS). Insight j, 2 (365):1–35.

Babiloni, C., Blinowska, K., Bonanni, L., Cichocki, A., De Haan, W., Del Percio, C., Dubois, B., Escudero, J., Fernández, A., Frisoni, G., Guntekin, B., Hajos, M., Hampel, H., Ifeachor, E., Kilborn, K., Kumar, S., Johnsen, K., Johannsson, M., Jeong, J., LeBeau, F., Lizio, R., Lopes da Silva, F., Maestú, F., McGeown, W. J., McKeith, I., Moretti, D. V., Nobili, F., Olichney, J., Onofrj, M., Palop, J. J., Rowan, M., Stocchi, F., Struzik, Z. M., Tanila, H., Teipel, S., Taylor, J. P., Weiergräber, M., Yener, G., Young-Pearse, T., Drinkenburg, W. H., and Randall, F. (2020). What electrophysiology tells us about Alzheimer’s disease: A window into the synchronization and connectivity of brain neurons. Neurobiology of Aging, 85:58–73.

Bao, Y., Yang, X., Fu, Y., Li, Z., Gong, R., and Lu, W. (2021). NMDAR-dependent somatic potentiation of synaptic inputs is correlated with β amyloid-mediated neuronal hyperactivity. Translational Neurodegeneration, 10(1):34.

Bastos, A. M., Usrey, W. M., Adams, R. A., Mangun, G. R., Fries, P., and Friston, K. J. (2012). Canonical microcircuits for predictive coding. Neuron, 76(4):695–711.

Beliveau, V., Ganz, M., Feng, L., Ozenne, B., L., H., M., F. P., C., S., N., G. D., and M., K. G. (2017). A high-resolution in vivo atlas of the human brain’s serotonin system. Journal of Neuroscience, 37(1):120–128.

Benwell, C. S. Y., Davila-Pérez, P., Fried, P. J., Jones, R. N., Travison, T. G., Santarnecchi, E., Pascual-Leone, A., and Shafi, M. M. (2020). EEG spectral power abnormalities and their relationship with cognitive dysfunction in patients with Alzheimer’s disease and type 2 diabetes. Neurobiology of Aging, 85:83–95.

Berger, M., Gray, J. A., and Roth, B. L. (2009). The expanded biology of serotonin. Annu Rev Med, 60:355–366.

Buzsáki, G., Anastassiou, C. A., and Koch, C. (2012). The origin of extracellular fields and currents — EEG, ECoG, LFP and spikes. Nature Reviews Neuroscience, 13(6):407–420.

Börgers, C., Epstein, S., and Kopell, N. J. (2008). Gamma oscillations mediate stimulus competition and attentional selection in a cortical network model. Proc Natl Acad Sci USA, 105(46):18023–18028.

Cameron, L. P. and Patel, S. D. e. a. (2023). 5-ht2ars mediate therapeutic behavioral effects of psychedelic tryptamines. ACS Chem Neurosci, 14(3):351–358.

Carhart-Harris, R. L. and Chandaria, S. e. a. (2023). Canalization and plasticity in psychopathology. Neuropharmacology, 226:109398.

Carhart-Harris, R. L. and Friston, K. J. (2019). REBUS and the anarchic brain: Toward a unified model of the brain action of psychedelics. Pharmacol Rev, 71(3):316–344.

Carhart-Harris, R. L. and Friston, K. J. (2019). REBUS and the anarchic brain: Toward a unified model of the brain action of psychedelics. Pharmacological Reviews, 71(3):316–344.

Carhart-Harris, R. L. and Leech, R. e. a. (2014). The entropic brain: a theory of conscious states informed by neuroimaging research with psychedelic drugs. Frontiers Human Neuroscience, 8:20.

Carhart-Harris, RL. (2018). The entropic brain — revisited. Neuropharmacology, 142(167–178).

Casula, E. P., Pellicciari, M. C., Bonnì, S., Borghi, I., Maiella, M., Assogna, M., Minei, M., Motta, C., D’Acunto, A., Porrazzini, F., Pezzopane, V., Mencarelli, L., Roncaioli, A., Rocchi, L., Spampinato, D. A., Caltagirone, C., Santarnecchi, E., Martorana, A., and Koch, G. (2022). Decreased Frontal Gamma Activity in Alzheimer Disease Patients. Annals of Neurology, 92(3):464.

Chetty, C. A., Bhardwaj, H., Kumar, G. P., Devanand, T., Sekhar, C. S. A., Aktürk, T., Kiyi, I., Yener, G., Güntekin, B., Joseph, J., and Adaikkan, C. (2024). EEG biomarkers in Alzheimer’s and prodromal Alzheimer’s: A comprehensive analysis of spectral and connectivity features. Alzheimer’s Research & Therapy, 16(1):236.

Cook, B. J., Peterson, A. D. H., Woldman, W., and Terry, J. R. (2022). Neural field models: A mathematical overview and unifying framework. Mathematical Neuroscience and Applications, 2.

Cover, T. M. (1999). Elements of information theory. John Wiley & Sons.

Deco, G. and Cruzat, J. e. a. (2018). Whole-brain multimodal neuroimaging model using serotonin receptor maps explains non-linear functional effects of LSD. Curr Biol, 28(19):3065–3074.

Desikan, R. S., Ségonne, F., Fischl, B., Quinn, B. T., Dickerson, B. C., Blacker, D., Buckner, R. L., Dale, A. M., Maguire, R. P., Hyman, B. T., et al. (2006). An automated labeling system for subdividing the human cerebral cortex on MRI scans into gyral based regions of interest. Neuroimage, 31(3):968–980.

Dhaynaut, M., Sprugnoli, G., Cappon, D., Macone, J., Sanchez, J. S., Normandin, M. D., Guehl, N. J., Koch, G., Paciorek, R., Connor, A., Press, D., Johnson, K., Pascual-Leone, A., El Fakhri, G., and Santarnecchi, E. (2022). Impact of 40 Hz Transcranial Alternating Current Stimulation on Cerebral Tau Burden in Patients with Alzheimer’s Disease: A Case Series. Journal of Alzheimer’s disease: JAD, 85(4):1667–1676.

Esteban, O., Markiewicz, C. J., Blair, R. W., Moodie, C. A., Isik, A. I., Erramuzpe, A., Kent, J. D., Goncalves, M., DuPre, E., Snyder, M., et al. (2019). fmriprep: a robust preprocessing pipeline for functional MRI. Nature methods, 16(1):111–116.

Fatemi, S. N., Sedghizadeh, M. J., and Aghajan, H. (2021). Theta-gamma phase-amplitude coupling explains the advantage of auditory plus visual gamma entrainment in Alzheimer’s therapy. Alzheimer’s & Dementia, 17(S7):e053451.

Fischl, B. (2012). Freesurfer. Neuroimage, 62(2):774–781.

Fischl, B., Van, D. K. A., Destrieux, C., Halgren, E., F., S., H., S. D., E., B., J., S. L., J., G., D., K., et al. (2004). Automatically parcellating the human cerebral cortex. Cerebral cortex, 14(1):11–22.

Friston, K. J., Mechelli, A., Turner, R., and Price, C. J. (2000). Non-linear responses in fmri: the balloon model, volterra kernels, and other hemodynamics. NeuroImage, 12(4):466–477.

Gallego-Rudolf, J., Wiesman, A. I., Pichet Binette, A., Villeneuve, S., and Baillet, S. (2024). Synergistic association of Aβ and tau pathology with cortical neurophysiology and cognitive decline in asymptomatic older adults. Nature Neuroscience, 27(11):2130–2137.

Garcés, P., Vicente, R., Wibral, M., Pineda Pardo, J. A., Lopez, M. E., Aurtenetxe, S., Marcos, A., de Andrés, M. E., Yus, M., Sancho, M., Maestú, F., and Fernández, A. (2013). Brain-wide slowing of spontaneous alpha rhythms in mild cognitive impairment. Frontiers in Aging Neuroscience, 5.

Garcia-Romeu, A., Darcy, S., Jackson, H., White, T., and Rosenberg, P. (2022). Psychedelics as Novel Therapeutics in Alzheimer’s Disease: Rationale and Potential Mechanisms. Current Topics in Behavioral Neurosciences, 56:287– 317.

Gaubert, S., Raimondo, F., Houot, M., Corsi, M.-C., Naccache, L., Diego Sitt, J., Hermann, B., Oudiette, D., Gagliardi, G., Habert, M.-O., Dubois, B., De Vico Fallani, F., Bakardjian, H., Epelbaum, S., and Alzheimer’s Disease Neuroimaging Initiative (2019). EEG evidence of compensatory mechanisms in preclinical Alzheimer’s disease. Brain, 142(7):2096–2112.

Gorgolewski, K., Burns, C. D., Madison, C., Clark, D., Halchenko, Y. O., Waskom, M. L., and Ghosh, S. S. (2011). Nipype: a flexible, lightweight and extensible neuroimaging data processing framework in python. Frontiers in neuroinformatics, 5:12318.

Halberstadt, A. L. (2015). Recent advances in the neuropsychopharmacology of serotonergic hallucinogens. Behavioural Brain Research, 277:99–120.

Hansen, J. Y., Shafiei, G., Markello, R. D., Smart, K., Cox, S. M. L., M., N., V., B., Y., W., J., G., É., A., et al. (2022). Mapping neurotransmitter systems to the structural and functional organization of the human neocortex. Nature neuroscience, 25(11):1569–1581.

Harris, S. S., Wolf, F., De Strooper, B., and Busche, M. A. (2020). Tipping the Scales: Peptide-Dependent Dysregulation of Neural Circuit Dynamics in Alzheimer’s Disease. Neuron, 107(3):417–435.

Herzog, R., Mediano, P. A. M., Rosas, F. E., Lodder, P., Carhart-Harris, R., Perl, Y. S., Tagliazucchi, E., and Cofre, R. (2023). A whole-brain model of the neural entropy increase elicited by psychedelic drugs. Scientific reports, 13(1):6244.

Husain, M. I., Ledwos, N., Fellows, E., Baer, J., Rosenblat, J. D., Blumberger, D. M., Mulsant, B. H., and Castle, D. J. (2023). Serotonergic psychedelics for depression: What do we know about neurobiological mechanisms of action? Frontiers Psychiatry, 13:1076459.

Iaccarino, H. F., Singer, A. C., Martorell, A. J., Rudenko, A., Gao, F., Gillingham, T. Z., Mathys, H., Seo, J., Kritskiy, O., Abdurrob, F., Adaikkan, C., Canter, R. G., Rueda, R., Brown, E. N., Boyden, E. S., and Tsai, L.-H. (2016). Gamma frequency entrainment attenuates amyloid load and modifies microglia. Nature, 540(7632):230–235.

Jansen, B. H. and Rit, V. G. (1995). Electroencephalogram and visual evoked potential generation in a mathematical model of coupled cortical columns. Biol Cybern, 73(4):357–366.

Jaramillo, J., Mejias, J. F., and Wang, X. (2019). Engagement of pulvino-cortical feedforward and feedback pathways in cognitive computations. Neuron, 101(2):321–336.

Jobst, B. M., Atasoy, S., Ponce-Alvarez, A., A., S., L., R., M., K., R., C.-H., L., K. M., and G., D. (2021). Increased sensitivity to strong perturbations in a whole-brain model of LSD. Neuroimage, 230:117809.

Jurcak, V., Tsuzuki, D., and Dan, I. (2007). 10/20, 10/10, and 10/5 systems revisited: Their validity as relative head-surface-based positioning systems. NeuroImage, 34(4):1600–1611.

Kometer, M., Schmidt, A., Jäncke, L., and Vollenweider, F. X. (2013). Activation of serotonin 2A receptors underlies the psilocybin-induced effects on α oscillations, N170 visual-evoked potentials, and visual hallucinations. The Journal of Neuroscience: The Official Journal of the Society for Neuroscience, 33(25):10544–10551.

Kozlowska, U., Nichols, C., Wiatr, K., and Figiel, M. (2022). From psychiatry to neurology: Psychedelics as prospective therapeutics for neurodegenerative disorders. Journal of Neurochemistry, 162(1):89–108.

Kringelbach, M. L. and Cruzat, J. e. a. (2020). Dynamic coupling of whole-brain neuronal and neurotransmitter systems. Proc Natl Acad Sci USA, 117(17):9566–9576.

López, M. E., Bruna, R., Aurtenetxe, S., Pineda-Pardo, J. Á., Marcos, A., Arrazola, J., Reinoso, A. I., Montejo, P., Bajo, R., and Maestú, F. (2014). Alpha-band hypersynchronization in progressive mild cognitive impairment: a magnetoencephalography study. Journal of Neuroscience, 34(44):14551–14559.

López-Sanz, D., Bruña, R., Garcés, P., Camara, C., Serrano, N., Rodríguez-Rojo, I. C., Delgado, M., Montenegro, M., López-Higes, R., Yus, M., et al. (2016). Alpha band disruption in the AD-continuum starts in the subjective cognitive decline stage: a MEG study. Scientific reports, 6(1):37685.

Luppi, A. I., Girn, M., Rosas, F. E., Timmermann, C., Roseman, L., Erritzoe, D., Nutt, D. J., Stamatakis, E. A., Spreng, R. N., Xing, L., et al. (2024). A role for the serotonin 2a receptor in the expansion and functioning of human transmodal cortex. Brain, 147(1):56–80.

Luppi, A. I., Mediano, P. A. M., Rosas, F. E., Allanson, J., Pickard, J. D., Williams, G. B., Craig, M. M., Finoia, P., Peattie, A. R. D., Coppola, P., et al. (2022). Whole-brain modelling identifies distinct but convergent paths to unconsciousness in anaesthesia and disorders of consciousness. Communications biology, 5(1):384.

Madsen, M. k., Fisher, P. M., Burmester, D., Dyssegaard, A., S., S. D., S., K., S., J. S., S., L., K., L., C., S., et al. (2019). Psychedelic effects of psilocybin correlate with serotonin 2a receptor occupancy and plasma psilocin levels. Neuropsychopharmacology, 44(7):1328–1334.

Markello, R. D., Hansen, J. Y., Liu, Z., Bazinet, V., Shafiei, G., E., S. L., N., B., J., S., S., B., D., S. T., et al. (2022). Neuromaps: structural and functional interpretation of brain maps. Nature Methods, 19(11):1472–1479.

Martorell, A. J., Paulson, A. L., Suk, H.-J., Abdurrob, F., Drummond, G. T., Guan, W., Young, J. Z., Kim, D. N.-W., Kritskiy, O., Barker, S. J., Mangena, V., Prince, S. M., Brown, E. N., Chung, K., Boyden, E. S., Singer, A. C., and Tsai, L.-H. (2019). Multi-sensory Gamma Stimulation Ameliorates Alzheimer’s-Associated Pathology and Improves Cognition. Cell, 177(2):256–271.e22.

Mejias, J. F., Murray, J. D., Kennedy, H., and Wang, X. (2016). Feed-forward and feedback frequency-dependent interactions in a large-scale laminar network of the primate cortex. Science advances, 2(11):e1601335.

Mercadal, B., Lopez-Sola, E., Galan-Gadea, A., Al, H. M., Sanchez-Todo, R., Salvador, R., Bartolomei, F., Wendling, F., and Ruffini, G. (2023). Towards a mesoscale physical modeling framework for stereotactic-EEG recordings. Journal of Neural Engineering, 20(1):016005.

Mindlin, I. and Herzog, R. e. a. (2023). Whole-brain modelling supports the use of serotonergic psychedelics for the treatment of disorders of consciousness. bioRxiv.

Miranda, PC., Mekonnen, A., Salvador, R., and Ruffini, G. (2013). The electric field in the cortex during transcranial current stimulation. Neuroimage, 70:48–58.

Müller, F., Lenz, C., Dolder, P., Lang, U., Schmidt, A., Liechti, M., and Borgwardt, S. (2017). Increased thalamic resting-state connectivity as a core driver of LSD-induced hallucinations. Acta Psychiatrica Scandinavica, 136(6):648–657.

Murty, D. V., Manikandan, K., Kumar, W. S., Ramesh, R. G., Purokayastha, S., Nagendra, B., ML, A., Balakrishnan, A., Javali, M., Rao, N. P., and Ray, S. (2021). Stimulus-induced gamma rhythms are weaker in human elderly with mild cognitive impairment and Alzheimer’s disease. eLife, 10:e61666. Publisher: eLife Sciences Publications, Ltd.

Muthukumaraswamy, S. D., Carhart-Harris, R. L., Moran, R. J., Brookes, M. J., Williams, T. M., Errtizoe, D., Sessa, B., Papadopoulos, A., Bolstridge, M., Singh, K. D., Feilding, A., Friston, K. J., and Nutt, D. J. (2013). Broadband Cortical Desynchronization Underlies the Human Psychedelic State. Journal of Neuroscience, 33(38):15171–15183.

Nakamura, A., Cuesta, P., Fernández, A., Arahata, Y., Iwata, K., Kuratsubo, I., Bundo, M., Hattori, H., Sakurai, T., Fukuda, K., Washimi, Y., Endo, H., Takeda, A., Diers, K., Bajo, R., Maestú, F., Ito, K., and Kato, T. (2018). Electromagnetic signatures of the preclinical and prodromal stages of Alzheimer’s disease. Brain: A Journal of Neurology, 141(5):1470–1485.

Nunez, P. L. and Srinivasan, R. (2006). Electric fields of the brain: the neurophysics of EEG. Oxford University Press, USA.

Nutt, D., Erritzoe, D., and Carhart-Harris, R. (2020). Psychedelic psychiatry’s brave new world. Cell, 181(1):24–29.

Ort, A., Smallridge, J. W., et al. (2023). TMS-EEG and resting-state EEG applied to altered states of consciousness: oscillations, complexity, and phenomenology. iScience, 26(5):106589.

Pallavicini, C., Vilas, M. G., Villarreal, M., Zamberlan, F., Muthukumaraswamy, S., Nutt, D., Carhart-Harris, R., and Tagliazucchi, E. (2019). Spectral signatures of serotonergic psychedelics and glutamatergic dissociatives. NeuroImage, 200:281–291.

Palop, J. J. and Mucke, L. (2016). Network abnormalities and interneuron dysfunction in Alzheimer disease. Nature Reviews. Neuroscience, 17(12):777– 792.

Passero, S., Rocchi, R., Vatti, G., Burgalassi, L., and Battistini, N. (1995). Quantitative EEG mapping, regional cerebral blood flow, and neuropsychological function in Alzheimer’s disease. Dementia (Basel, Switzerland), 6(3):148–156.

Petersen, R. C. (2004). Mild cognitive impairment as a diagnostic entity. Journal of Internal Medicine, 256(3):183–194.

Press, W. H. and Teukolsky, S. A. (1992). Adaptive stepsize runge-kutta integration. Comput Phys, 6(2):111–216.

Puttaert, D., Wens, V., Fery, P., Rovai, A., Trotta, N., Coquelet, N., De Breucker, S., Sadeghi, N., Coolen, T., Goldman, S., Peigneux, P., Bier, J.-C., and De, Tiège, X. (2021). Decreased Alpha Peak Frequency Is Linked to Episodic Memory Impairment in Pathological Aging. Frontiers in Aging Neuroscience, 13.

Rochart, R., Liu, Q., Fonteh, A. N., Harrington, M. G., and Arakaki, X. (2020). Compromised behavior and gamma power during working memory in cognitively healthy individuals with abnormal CSF amyloid/tau. Frontiers in Aging Neuroscience, 12:574214.

Ronneberger, O., Fischer, P., and Brox, T. (2015). U-net: Convolutional networks for biomedical image segmentation. In Navab, N., Hornegger, J., Wells, W. M., and Frangi, A. F., editors, Medical Image Computing and Computer-Assisted Intervention – MICCAI 2015, pages 234–241, Cham. Springer International Publishing.

Ruffini, G. (2015). Application of the reciprocity theorem to EEG inversion and optimization of EEG-driven transcranial current stimulation (tcs, including tdcs, tacs, trns). arXiv.

Ruffini, G. (2017). Lempel-Zip Complexity Reference.

Ruffini, G., Castaldo, F., Lopez-Sola, E., Sanchez-Todo, R., and Vohryzek, J. (2024a). The algorithmic agent perspective and computational neuropsychiatry: From etiology to advanced therapy in major depressive disorder. PsyArXiv.

Ruffini, G., Damiani, G., Lozano-Soldevilla, D., Deco, N., Rosas, F. E., Kiani, N. A., Ponce-Alvarez, A., Kringelbach, M. L., Carhart-Harris, R., and Deco, G. (2023). LSD-induced increase of Ising temperature and algorithmic complexity of brain dynamics. Plos Computational Biology, 19(2):e1010811.

Ruffini, G., Fox, M. D., Ripolles, O., Miranda, P. C., and Pascual-Leone, A. (2014). Optimization of multifocal transcranial current stimulation for weighted cortical pattern targeting from realistic modeling of electric fields. Neuroimage, 89:216–225.

Ruffini, G., Ibañez, D., Kroupi, E., Gagnon, J.-F., Montplaisir, J., Postuma, R. B., Castellano, M., and Soria-Frisch, A. (2019). Algorithmic complexity of EEG for prognosis of neurodegeneration in idiopathic rapid eye movement behavior disorder (RBD). Annals of biomedical engineering, 47:282–296.

Ruffini, G., Lopez-Sola, E., Palma, R., Vohryzek, J., Castaldo, F., and Friston, K. (2025). Cross-frequency coupling as a neural substrate for prediction error evaluation: A laminar neural mass modeling approach. bioRxiv, pages 2025–03.

Ruffini, G., Lopez-Sola, E., Vohryzek, J., and Sanchez-Todo, R. (2024b). Neural geometrodynamics, complexity, and plasticity: a psychedelics perspective. Entropy, 26(1).

Ruffini, G., Sanchez-Todo, R., Dubreuil, L., Salvador, R., Pinotsis, D., Miller, E. K., Wendling, F., E. Santarnecchi, and Bastos, A. (2020). P118 A Bio-physically realistic Laminar Neural Mass Modeling framework for transcranial Current Stimulation. Clinical Neurophysiology, 131(4):e78–e79.

Ruffini, G., Wendling, F., Merlet, I., Molaee-Ardekani, B., Mekonnen, A., Salvador, R., Soria-Frisch, A., Grau, C., Dunne, S., and Miranda, P. C. (2013). Transcranial Current Brain Stimulation (tCS): Models and Technologies. IEEE Transactions on Neural Systems and Rehabilitation Engineering, 21(3):333–345.

Salvador, R., Biagi, M., Pelegrí, M. P., Zhou, J., Travison, T., Pascual-Leone, A., Manor, B., and Ruffini, G. (2022). Towards the identification and optimization of the “dose-response” relationship of transcranial direct current stimulation. bioRxiv, pages 2022–01.

Sanchez-Todo, R. and Bastos, A. M. e. a. (2023). A physical neural mass model framework for the analysis of oscillatory generators from laminar electrophysiological recordings. Neuroimage, 270:119938.

Sanchez-Todo, R., Lopez-Sola, E., Mercadal, B., and Ruffini, G. (2024). Laminar Neural Mass Model for Representing Alzheimer’s Disease Electrophysiopathology. in prep.

Sanchez-Todo, R., Salvador, R., Santarnecchi, E., Wendling, F., Deco, G., and Ruffini, G. (2018). Personalization of hybrid brain models from neuroimaging and electrophysiology data. bioRxiv.

Silverstein, B. H., Kolbman, N., Nelson, A., Liu, T., Guzzo, P., Gilligan, J., Lee, U., Mashour, G. A., Vanini, G., and Pal, D. (2024). Psilocybin induces dose-dependent changes in functional network organization in rat cortex.

Smausz, R., Neill, J., and Gigg, J. (2022). Neural mechanisms underlying psilocybin’s therapeutic potential – the need for preclinical in vivo electrophysiology. Journal of Psychopharmacology, 36(7):781–793.

Sperling, R. A., Aisen, P. S., Beckett, L. A., Bennett, D. A., Craft, S., Fagan, A. M., Iwatsubo, T., Jack, C. R., Kaye, J., Montine, T. J., Park, D. C., Reiman, E. M., Rowe, C. C., Siemers, E., Stern, Y., Yaffe, K., Carrillo, M. C., Thies, B., Morrison-Bogorad, M., Wagster, M. V., and Phelps, C. H. (2011). Toward defining the preclinical stages of Alzheimer’s disease: Recommendations from the National Institute on Aging-Alzheimer’s Association workgroups on diagnostic guidelines for Alzheimer’s disease. Alzheimer’s & Dementia: The Journal of the Alzheimer’s Association, 7(3):280– 292.

Sprugnoli, G., Munsch, F., Cappon, D., Paciorek, R., Macone, J., Connor, A., El Fakhri, G., Salvador, R., Ruffini, G., Donohoe, K., Shafi, M. M., Press, D., Alsop, D. C., Pascual Leone, A., and Santarnecchi, E. (2021). Impact of multisession 40Hz tACS on hippocampal perfusion in patients with Alzheimer’s disease. Alzheimer’s Research & Therapy, 13(1):203.

Sun, J., Wang, B., Niu, Y., Tan, Y., Fan, C., Zhang, N., Xue, J., Wei, J., and Xiang, J. (2020). Complexity Analysis of EEG, MEG, and fMRI in Mild Cognitive Impairment and Alzheimer’s Disease: A Review. Entropy, 22(2):239.

Tagliazucchi, E., Roseman, L., Kaelen, M., Orban, C., Muthukumaraswamy, S. D., Murphy, K., Laufs, H., Leech, R., McGonigle, J., Crossley, N., Bullmore, E., Williams, T., Bolstridge, M., Feilding, A., Nutt, D. J., and Carhart-Harris, R. (2016). Increased Global Functional Connectivity Correlates with LSD-Induced Ego Dissolution. Current biology: CB, 26(8):1043–1050.

Timmermann, C., Roseman, L., et al. (2023). Human brain effects of DMT assessed via EEG-fmri. Proc Natl Acad Sci USA, 120(13):e2218949120.

Tournier, J.-D., Smith, R., Raffelt, D., Tabbara, R., Dhollander, T., Pietsch, M., Christiaens, D., Jeurissen, B., Yeh, C.-H., and Connelly, A. (2019). Mrtrix3: A fast, flexible and open software framework for medical image processing and visualisation. Neuroimage, 202:116–137.

Uddin, L. Q., Mooshagian, E., Zaidel, E., Scheres, A., Margulies, D. S., Kelly, A. M. C., Shehzad, Z., Adelstein, J. S., Castellanos, F. X., Biswal, B. B., et al. (2008). Residual functional connectivity in the split-brain revealed with resting-state functional MRI. Neuroreport, 19(7):703–709.

Vann Jones, S. A. and O’Kelly, A. (2020). Psychedelics as a Treatment for Alzheimer’s Disease Dementia. Frontiers in Synaptic Neuroscience, 12:34.

Vohryzek, J., Cabral, J., Lord, L., Fernandes, H., Roseman, L., Nutt, D., Carhart-Harris, R., Deco, G., and Kringelbach, M. (2024). Brain dynamics predictive of response to psilocybin for treatment-resistant depression. Brain Communications, 6(2):fcae049.

Vohryzek, J., Cabral, J., Vuust, P., Deco, G., and Kringelbach, M. (2022). Understanding brain states across spacetime informed by whole-brain modelling. Philosophical Transactions of the Royal Society A, 380(2227):20210247.

Wang, R., Wang, J., Yu, H., Wei, X., Yang, C., and Deng, B. (2015). Power spectral density and coherence analysis of Alzheimer’s EEG. Cognitive Neurodynamics, 9(3):291–304.

Wang, X., Leong, A. T. L., Chan, R. W., Liu, Y., and Wu, E. X. (2019). Thalamic low frequency activity facilitates resting-state cortical interhemispheric MRI functional connectivity. Neuroimage, 201:115985.

Warren, A. L., Lankri, D., Cunningham, M. J., Serrano, I. C., Parise, L. F., Kruegel, A. C., Duggan, P., Zilberg, G., Capper, M. J., Havel, V., et al. (2024). Structural pharmacology and therapeutic potential of 5-methoxytryptamines. Nature, pages 1–10.

Welch, P. (1967). The use of fast fourier transform for the estimation of power spectra: A method based on time averaging over short, modified periodograms. IEEE Transactions on Audio and Electroacoustics, 15(2):70–73.

Welch, T. A. (1984). A technique for high-performance data compression. Computer, 17(06):8–19.

Winkelman, M. J., Szabo, A., and Frecska, E. (2023). The potential of psychedelics for the treatment of Alzheimer’s disease and related dementias. European Neuropsychopharmacology: The Journal of the European College of Neuropsychopharmacology, 76:3–16.

Zott, B., Simon, M. M., Hong, W., Unger, F., Chen-Engerer, H.-J., Frosch, M. P., Sakmann, B., Walsh, D. M., and Konnerth, A. (2019). A vicious cycle of β amyloid-dependent neuronal hyperactivation. Science (New York, N.Y.), 365(6453):559– 565.

